# Aromatic polyketide biosynthesis: fidelity, evolution and engineering

**DOI:** 10.1101/581074

**Authors:** Zhiwei Qin, Rebecca Devine, Matthew I. Hutchings, Barrie Wilkinson

## Abstract

We report the formicapyridines which are structurally and biosynthetically related to the pentacyclic fasamycin and formicamycin aromatic polyketides but comprise a rare pyridine moiety. These new compounds are trace level metabolites formed by derailment of the major biosynthetic pathway. Inspired by evolutionary logic we show that rational mutation of a single gene in the biosynthetic gene cluster leads to a significant increase both in total formicapyridine production and their enrichment relative to the fasamycins/formicamycins. Our observations broaden the polyketide biosynthetic landscape and identify a non-catalytic role for ABM superfamily proteins in type II polyketide synthase assemblages for maintaining biosynthetic pathway fidelity.

## Introduction

An idealised linear biosynthetic pathway to a complex natural product can be imagined proceeding through a series of intermediate structures which would exist for some finite time as the pathway product accumulates. These hypothetical intermediates would exist freely in solution or bound to enzymes in the case of assembly line processes, and eventually all flux through the pathway would end and only the final product would exist. In reality no pathways proceed in this manner as the situation is complicated by varying rates of reaction for the different steps, meaning that some intermediates accumulate at significant concentrations, while the inherent reactivity of other intermediates, or their ability to act as substrates for housekeeping enzymes not dedicated to the pathway, means that shunt metabolites often arise. Matters are further complicated by the fact that pathways are often convergent, with multiple units made in parallel before assembly into the final product, for example in the biosynthetic pathways to macrolide or aminoglycoside^1^ antibiotics. In addition, some pathways are not linear, and the final product is accessed *via* several routes, due to the inherent substrate plasticity of the biosynthetic enzymes; well-studied examples include the rapamycin and erythromycin pathways^2^. Thus, in practice, any biosynthetic pathway will lead to the accumulation of a mixture comprising the final product plus varying concentrations of pathway intermediates and shunt metabolites. The composition of such a mixture will vary further when alternate growth conditions are used^3^. Sometimes the ‘final’ product of the pathway cannot even be clearly discerned.

Such mixtures of compounds are said to comprise a series of biosynthetic congeners, and their identification can provide valuable information about a biosynthetic pathway and may sometimes lead to new biosynthetic understanding^4^. We recently reported the formicamycins, pentacyclic polyketides produced by *Streptomyces formicae* KY5 isolated from the fungus-growing plant-ant *Tetroponera penzigi* (Fig. 1).^5^ In total, sixteen congeners were isolated during the initial study, including three new fasamycins, a group of compounds previously reported from the heterologous expression of a clone derived from environmental DNA^6^. The fasamycins are likely to be biosynthetic precursors of the formicamycins. These various congeners are the product of a type II polyketide synthase (PKS) operating in conjunction with a series of post-PKS modifications that include O-methylation and halogenation, plus oxidative and reductive modifications. Intrigued by these compounds, which exhibit potent antibacterial activity, and as part of studies to decipher their biosynthetic pathway, we employed targeted metabolomics to identify further congeners that may have been missed during manual analysis of culture extracts. This led us to identify the formicapyridines (**1-9**), pyridine containing polyketide alkaloids which represent additional products of the formicamycin *(for)* biosynthetic gene cluster (BGC). Products of type II PKS systems which contain a pyridine moiety are extremely rare^7^.

**Figure 1.**
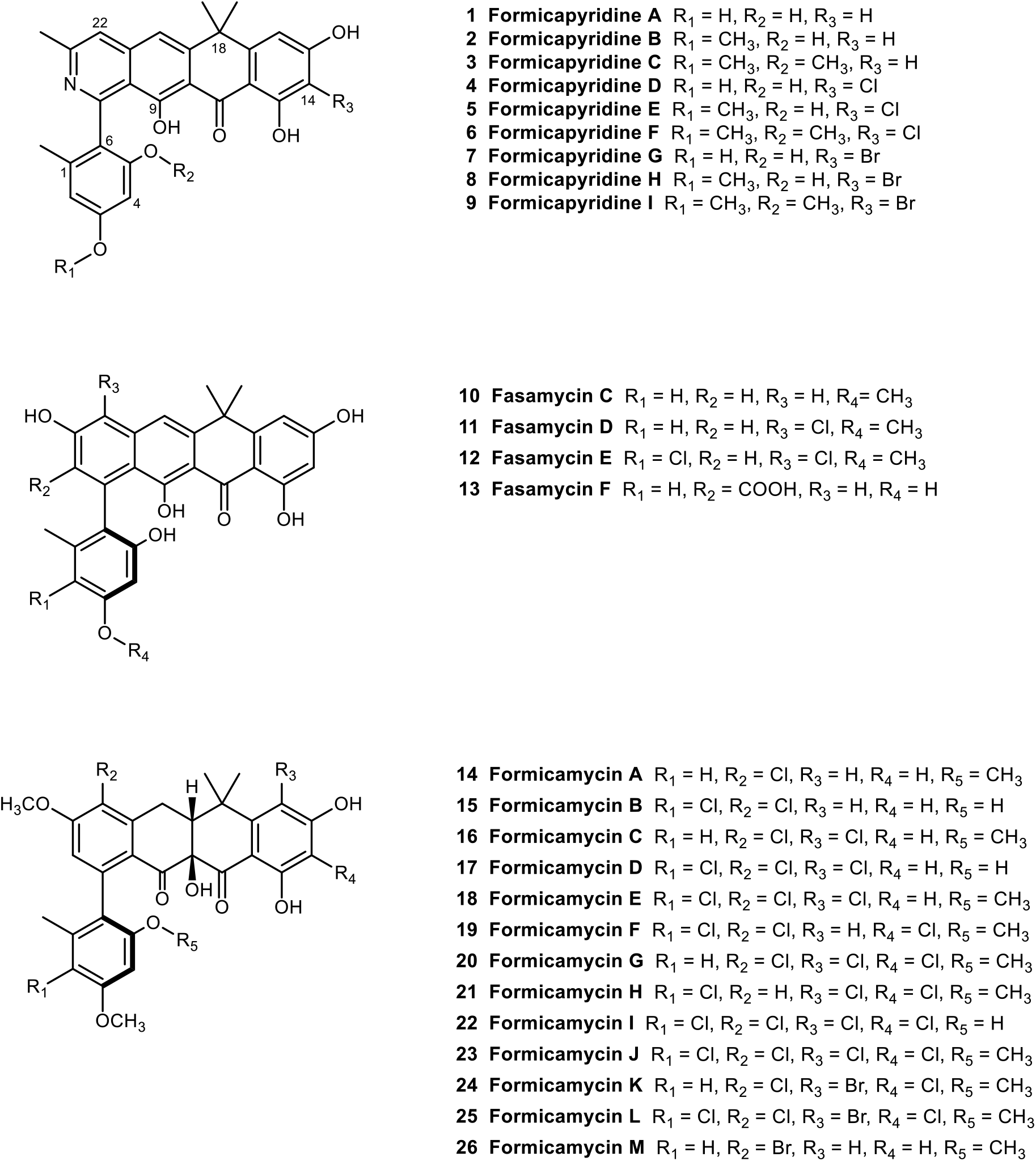
Chemical structures of metabolites isolated from *Streptomyces formicae*. Compounds **1-9** and **13** were discovered during this study whereas **10-12** and **14-26** were reported in an earlier study^5^.

Challis and co-workers recently showed that the majority of actinomycete derived polyketide alkaloids, including those containing a pyridine moiety, arise from reactive intermediates formed after transamination of aldehydes generated from reductive off-loading of the thioester bound polyketide chain from a type I modular PK^8^. In contrast, the formicapyridines are minor shunt metabolites that likely arise due to derailment of the formicamycin biosynthetic pathway. Intrigued by these observations we investigated the possibility of reprogramming, or evolving, the BGC such that the formicapyridines might become the major products of the formicamycin BGC. Following bioinformatic and mutational analysis we identified a mutant Δ*forS* which significantly increased the production of formicapyridines while reducing the combined titre of fasamycins and formicamycins. These and other mutational data lead us to the hypothesis that ForS is not a cyclase but forms part of a multienzyme complex where it acts as a chaperone-like protein to aid in maintaining pathway fidelity and performance. The discovery and engineering of formicapyridine biosynthesis raises intriguing questions regarding the evolution of type II PKS biosynthetic pathways and the origins of natural product chemical diversity.

## Results

### Metabolomics led identification of the formicapyridines

We attempted to identify new biosynthetic congeners using molecular networking via the Global Natural Product Social Molecular Networking (GNPS) web-platform^9^. However, the aromatic, polycyclic nature of these molecules limited fragmentation and the effectiveness of GNPS. Instead, as most fasamycin and formicamycin congeners are halogenated, we established a bespoke dereplication method by making use of *S. formicae* KY5 mutants that we reported previously, including an entire BGC deletion strain (Δ*for*), and a strain in which the pathway specific halogenase gene was deleted (Δ*forV*)^5^ (Fig. 2). Our earlier study^5^ showed that the Δ*for* mutant does not produce any fasamycins or formicamycins, and the Δ*forV* mutant produces only non-halogenated fasamycin congeners.

**Figure 2.**
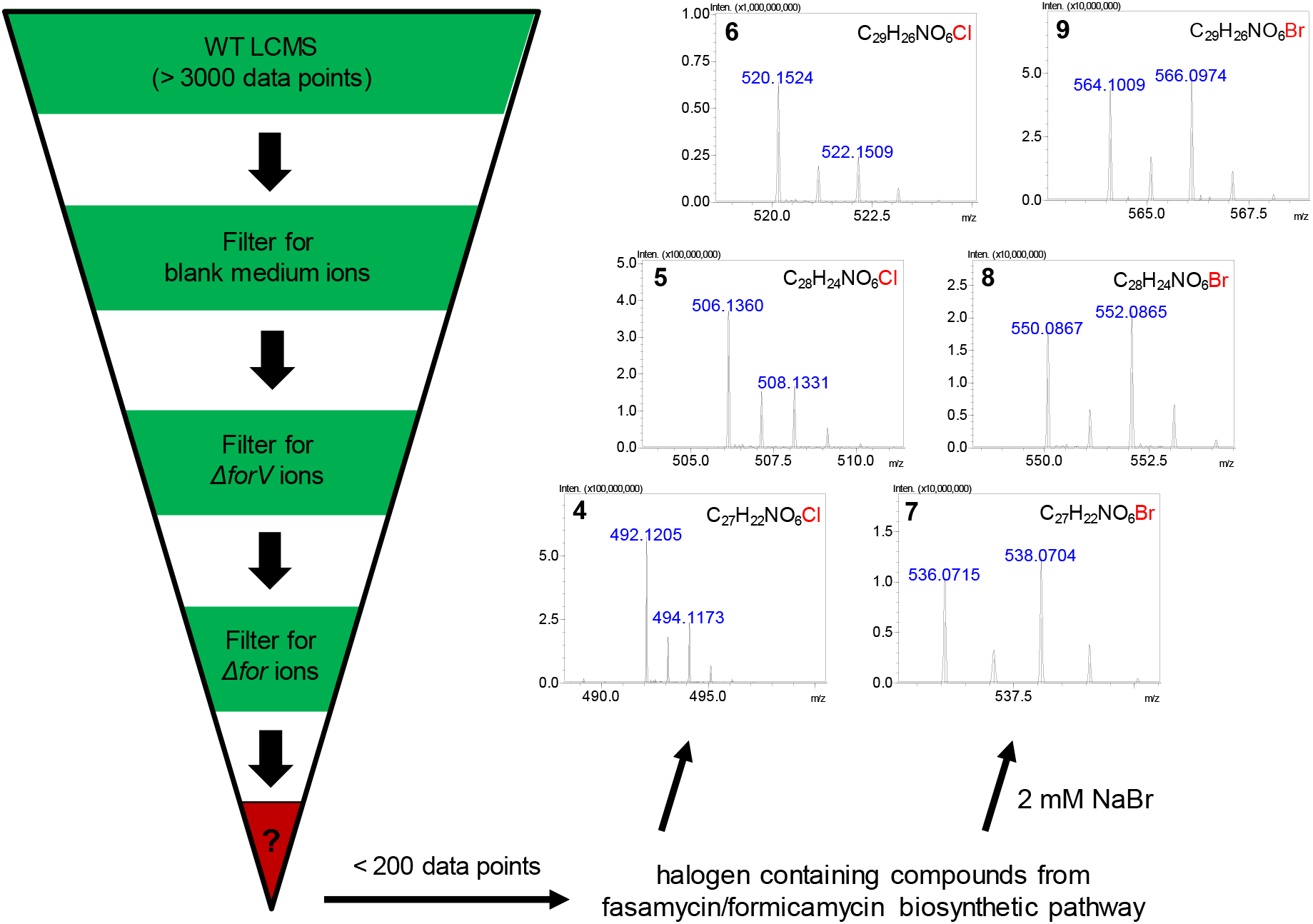
Metabolomics pipeline. Dereplication, based on *for* biosynthetic mutants and exogenous bromide addition, led to the identification of the new congeners **4-9** with characteristic halogen-containing patterns (strain Δ*forV* lacks a halogenase gene; Δ*for* lacks the entire biosynthetic gene cluster).

Replicate (n=3) ethyl acetate extracts of the wild type (WT), Δ*for* and Δ*forV* strains, along with equivalent extracts from uninoculated mannitol soya flour (MS) agar plates, were analysed by liquid chromatography high-resolution mass spectroscopy (LC-HRMS). Using the Profiling Solutions software (Shimadzu Corporation), the WT dataset (Supplementary Dataset 1) was filtered to remove ions also present in the other samples. Only two other BGCs in the *S. formicae* KY5 genome^10^ contain putative halogenase encoding genes, so we hypothesised that any chlorine-containing ions present in the filtered dataset would likely derive from the formicamycin biosynthetic pathway. This process dramatically reduced the dataset complexity, leaving less than 200 unique ions from an original set of approx. 3000. Manual curation showed that most of the remaining ions corresponded to halogenated molecules (based on isotope patterns), leaving twelve previously identified fasamycin and formicamycin congeners, two new fasamycin/formicamycin congeners that remain uncharacterized due to trace levels, and the known isoflavone 6-chlorogenistein^11^ along with a regioisomer (Supplementary Note 1). Analysis of C/H ratios and *m/z* data allowed us to identify a group of three additional new metabolites (**4-6**) with mass spectra suggesting a close structural relationship to the fasamycins/formicamycins; these varied only in the number of methyl groups present. By searching for equivalent ions lacking chlorine atoms we identified three additional congeners (**1-3**). Compounds **1-6** were not initially observed in the UV chromatograms of the WT, and qualitative examination of the LCMS data suggested titres at least 100-fold lower than for the formicamycins. Furthermore, and surprisingly, the *m/z* data implied the presence of a single nitrogen atom.

We then repeated the experiment but included a set of WT strain fermentations to which sodium bromide (2 mM) was added, as our previous study showed this leads to the biosynthesis of brominated formicamycin congeners^5^ This allowed the identification of three conditional bromine-containing metabolites (**7-9**) with MS characteristics like those of **4-6** (Fig. 2 and Supplementary Figure 1), lending support to the hypothesis that these compounds represent a family of biosynthetic congeners.

### Isolation, structure elucidation and biological activity

HRMS data allowed us to predict molecular formulae for **1-9** that were used to search the online chemical database REAXYS^12^. This suggested that they represented new structures. To isolate sufficient material for structure determination and antibacterial assays, the growth of *S. formicae* KY5 was scaled up (14 L; ~450 MS agar plates). After nine days incubation at 30°C the combined agar was chopped up, extracted with ethyl acetate, and the solvent removed under reduced pressure. The resulting extract was subjected to repeated rounds of reversed-phase HPLC followed by Sephadex LH-20 size exclusion chromatography. In this way, small quantities of purified **1-6** were isolated.

We first determined the structure of **2** which was isolated in the largest amount (~2 mg). HRMS and ^13^C NMR data indicated a probable molecular formula of C_28_H_25_NO_6_ (calculated *m/z* 472.1755 ([M+H]^+^); observed *m/z* 472.1753 ([M+H]^+^); Δ = −0.42 ppm) indicating 17 degrees of unsaturation. The UV spectrum showed absorption maxima at 229, 249, 272 and 392 nm indicating a complex conjugated system that was somewhat different to that of the fasamycins and formicamycins.^5^ Inspection of the ^13^C NMR spectrum showed 21 sp^2^ carbons (δ_C_ 99.90-167.54 ppm), one carbonyl carbon (δ_C_ 191.81 ppm), four methyl carbons (δ_C_ 20.41, 23.59, 34.56 and 37.75 ppm), one methoxy carbon (δ_C_ 55.80 ppm) and one sp^3^ quaternary carbon (δ_C_ 40.49 ppm). The ^1^H NMR spectrum revealed the presence of five methyl singlets (δ_H_ 1.75, 1.76, 1.93, 3.67 and 3.81 ppm), two aromatic proton singlets (δ_H_ 7.62 and 7.65 ppm), plus four aromatic proton doublets (δ_H_ 6.24 (d, 2.25 Hz), 6.70 (d, 2.25 Hz), 6.34 (d, 2.27 Hz) and 6.42 (d, 2.27 Hz)). Limited ^1^H-^1^H COSY correlations meant we were reliant upon HMBC-based atomic connections, and through-space NOESY correlations, which led to two potential substructures consisting of 26 carbon atoms, leaving one carbonyl and one phenol carbon unassigned (Fig. 3). This left some uncertainty, but given the relationship with the fasamycins, we predicted the structure to be that shown for **2**. We thus plotted the ^13^C chemical shifts for **2** against those for fasamycin C (**10**), the closest structural congener, and then all other congeners. The data were in excellent agreement with the main differences at C7, C22 and C23, consistent with the adjacent nitrogen atom (Supplementary Figure 2). With the structure of **2** in hand we could readily assign the structures of **1** and **3-6** (Supplementary Note 2). NOESY correlations played a key role allowing us to link the methoxy group at C3 with H2 and H4 (e.g. **2, 3, 5** and **6**), and the methoxy at C5 with H4 (e.g. **3** and **6**). We also used NOESY and HSQC correlations to distinguish C14 and C16 once one was chlorinated: given the NOESY correlations to the *gem*-dimethyl groups of C26 and C27 with H16, and the disappearance of a HSQC linkage for C14, we concluded that C14 was chlorinated (e.g. **4-6**). Compounds **1-6** exhibit optical activity with small [α]_D_^20^ values between +8° and +13°, indicating a preferred conformation about the chiral axis of the C6-C7 bond. However, we were unable to assign the stereochemistry due to the small amounts of compound and very weak electronic circular dichroism (ECD) spectra. Due to the very low levels of production, we have assigned preliminary structures for **7-9** based on **4-6**, with a bromine atom replacing chlorine at C14.

**Figure 3.**
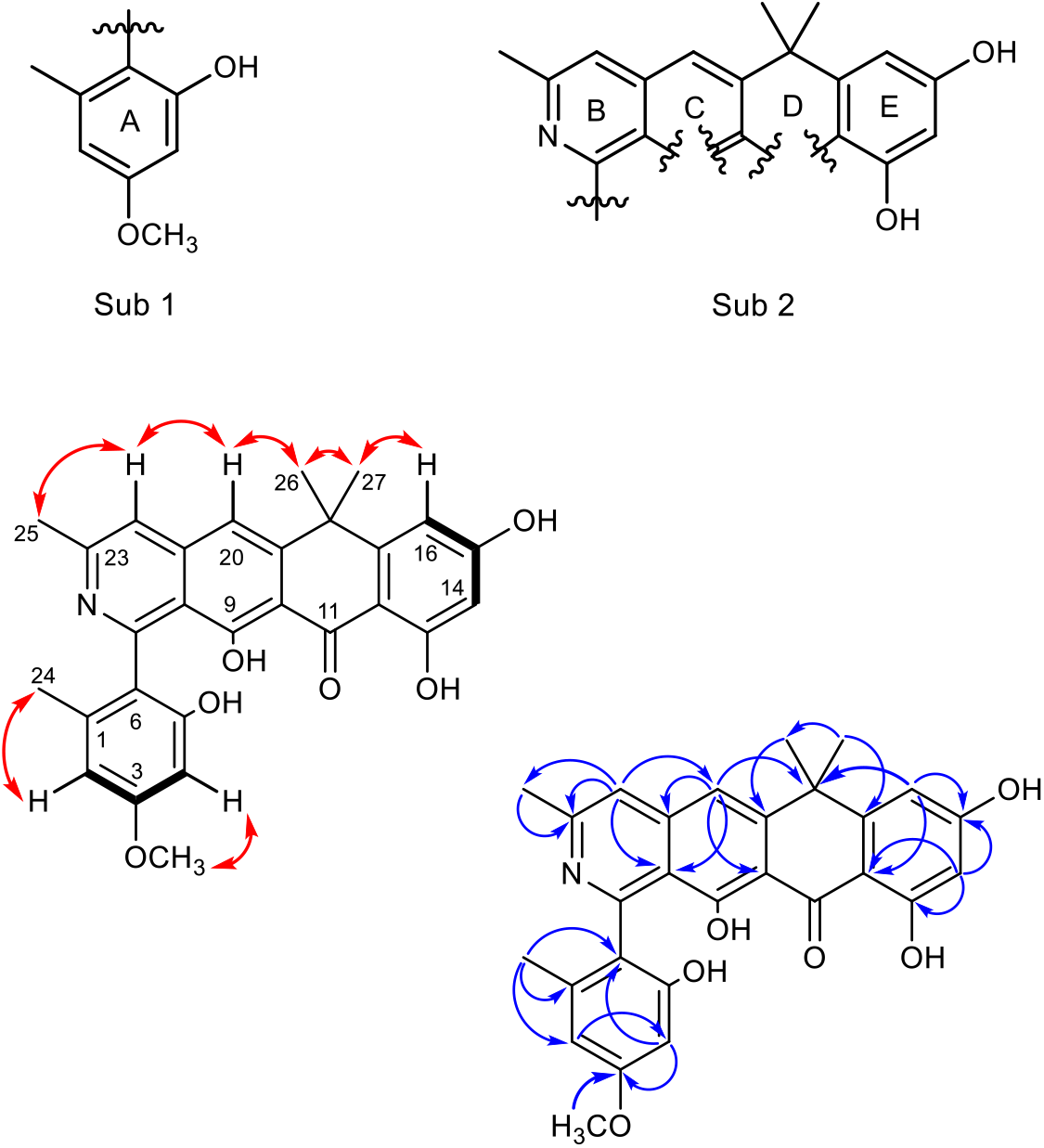
Structure determination. The COSY (black bold), NOSEY (red double head arrows) and HMBC (blue single headed arrows) correlations for formicapyridine B (**2**), along with the two resulting substructures are shown (Sub 1 and Sub 2).

Compounds **1-6** displayed no antibacterial activity against *Bacillus subtilis* 168^13^ using overlay assays at 5 μg/mL, the average concentration used for illustrating fasamycin and formicamycin bioactivity. Similarly, no inhibition was seen at ten-fold this concentration (50 μg/mL), with only small zones of inhibition at 100-fold (500 μg/mL). To confirm this was not a result of reduced diffusion from the disc, assays were set up to test growth of *B. subtillis* in liquid cultures containing compounds **1-6**. Again, no significant reduction in colony forming units (CFU/mL) was seen.

### Biosynthetic origins

Interrogation of LC-HRMS data from extracts of the Δ*for* and Δ*forV* mutants verified that the fasamycin/formicamycin biosynthetic machinery is required for formicapyridine production. The Δ*for* mutant does not produce **1-6** (Supplementary Figure 3b), but the production of all six compounds was restored upon ectopic expression of the P1-derived artificial chromosome (PAC) pESAC13-215-G, which contains the entire formicamycin BGC (Supplementary Figure 3c). Similarly, the Δ*forV* mutant does not produce **4-6** (Supplementary Figure 3d), but their production is re-established upon ectopic expression of *forV* under the control of the native promoter using an integrative plasmid (Supplementary Figure 3e) (the construction of all strains, the requirement of the *for* BGC for production of the fasamycins and formicamycins, and the requirement of ForV for their halogenation were described previously^5^.)

The very low level of formicapyridines made by the WT strain suggests they are shunt metabolites arising from aberrant derailment of fasamycin/formicamycin biosynthesis. On this basis, we suggest a biosynthetic pathway as described in Fig. 4. Assembly of the poly-β-ketone tridecaketide intermediate should proceed as previously proposed on route to the fasamycins^5^. This would be followed by a series of cyclization and aromatisation steps, presumably in a sequential manner, with the final cyclization event probably involving formation of the B-ring. However, premature hydrolysis of the acylcarrier protein from putative intermediate **27**, prior to the action of a final cyclase, would liberate the enzyme-free, β-ketoacid species **28** that would be highly facile to spontaneous decarboxylation yielding the methylketone **29**. An endogenous aminotransferase from the cellular milieu would then generate either of the species **30** or **31**, which could undergo cyclization, dehydration and oxidation to yield formicapyridine **1**.

**Figure 4.**
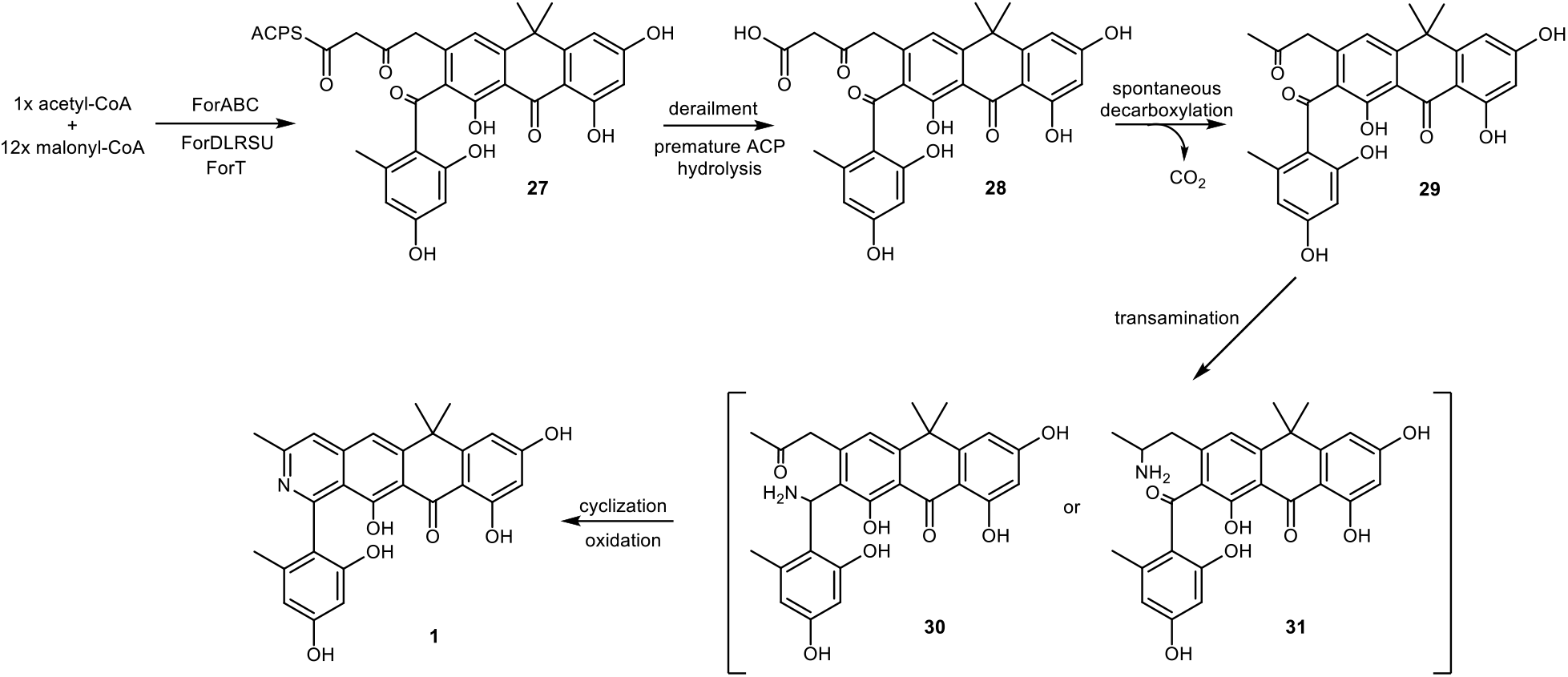
Proposed biosynthesis of the formicapyridine backbone. This pathway requires that C25 of the formicapyridines originates from C2 of an acetate unit, with C1 lost via decarboxylation. To support our hypothesis, and the backbone structural assignments, *S. formicae* KY5 was cultivated on MS agar plates for two days and then overlaid with a solution of [1,2-^13^C_2_] sodium acetate (1 mL of 60 mM solution; final concentration 2 mM). This was repeated on the four following days, and after a total of 9 days incubation the agar was extracted with ethyl acetate and the most abundant congener was isolated (**2**, ~1 mg), and the ^13^C NMR spectrum was acquired. Due to the small amount of material, and overlapping signals, only eight of the intact acetate units could be unambiguously identified based on their coupling contests in addition to an enriched singlet for C25 (Fig. 5; Supplementary Figure 4). This is consistent with our biosynthetic hypothesis which requires that the carbon atom at C25 derives from C2 of a fragmented acetate unit. The data were in accordance with the proposed structure for **2**.

### Targeted evolution of the *for* BGC

The biosynthetic proposal above led us to the following thought experiment. Suppose, in some environmental scenario, the presence of formicapyridines leads to a selection advantage. Is there then a single mutation in the BGC that could rapidly lead to significantly enhanced levels of their production, and, in addition, could such changes lead to the reduction or even abolition of fasamycin/formicamycin biosynthesis?

The proposed fasamycin biosynthetic pathway likely requires the action of multiple polyketide cyclase/aromatase enzymes and this suggests that one cyclase could be dedicated to formation of ring-B in the final step of backbone biosynthesis (Fig. 4). On this basis we hypothesised that mutation of a gene encoding a putative ring-B cyclase might lead to the phenotype imagined in our thought experiment by eliminating the biosynthesis of fasamycins/formicamycins and shunting carbon flux into the proposed formicapyridine pathway. As described below, bioinformatics (BLAST and conserved domain analysis), in conjunction with structural modelling using the Phyre2 web portal for protein modelling and analysis^14^, allowed us to identify five genes in the *for* BGC which might encode potential cyclase/aromatase enzymes (Table 1).

**Table 1.** Characteristics of the putative PKS cyclase genes in the for BGC.

| Name | Sequence identity | Top BLASTp hit | Annotation | % coverage | % identity |
| --- | --- | --- | --- | --- | --- |
| **ForD** | WP_098245758.1 | WP_003962252.1 | Polyketide cyclase (*TcmN*), *Streptomyces clavuligerus* ATCC 27064 | 95 | 73 |
| **ForL** | WP_055544278.1 | WP_076971949.1 | *Tcml* family type II polyketide cyclase, *Streptomyces sparsogenes* | 97 | 52 |
| **ForR** | WP_098245762.1 | AUI41024.1 | Polyketide WhiE II *Streptomyces* sp., cupin domain | 96 | 75 |
| **ForS** | WP_098245763 | SDF47214.1 | Antibiotic biosynthesis monooxygenase, *Lechevalieria fradiae* | 96 | 48 |
| **ForU** | WP_098245765.1 | PZS28546.1 | Antibiotic biosynthesis monooxygenase, *Pseudonocardiales* bacteria | 84 | 46 |

The gene product ForD shows significant sequence similarity to aromatase/cyclases (ARO/CYC) such as the N-terminal domain (pfam 03364) of the archetypical tetracenomycin polyketide cyclase TcmN which belongs to the Bet v1-like superfamily (cl10022)^15^. Extensive *in vivo* analysis^16, 17, 18^ and *in vitro* reconstruction^19^ showed that TcmN catalyses formation of the first two rings of tetracenomycin *via* sequential C9-C14 and C7-C16 cyclization/aromatization reactions. Thus, we predict ForD will play a key role in formation of the rings E and D during fasamycin biosynthesis. The ForL gene product belongs to the TcmI family of polyketide cyclases (cl24023; pfam 04673). The function of TcmI has also been verified by *in vivo* mutational analysis^20^ and biochemical characterisation^21, 22^. It catalyses cyclization of the final ring during formation of tetracenomycin F1 from tetracenomycin F2, and this reaction is remarkably like that proposed in our hypothesis for the formation of ring-B as the final step of fasamycin backbone assembly (Fig. 4). ForR is a homologue of the zinc containing polyketide cyclase RemF from the resistomycin BGC^23, 24^. RemF is a single domain protein (pfam 07883) comprising the conserved barrel domain of the cupin superfamily (cl21464)^25^. Finally, the gene products ForS and ForU are both single domain proteins (pfam 03992) belonging to the ABM superfamily (cl10022). A notable member of this family is ActVA-orf6 which functions as a monooxygenase during biosynthesis of the polyketide antibiotic actinorhodin; the structure of this enzyme has been solved and it is topologically related to the PKS cyclase TcmI and homologues^26^. Notably, ABM family domains are found in several PKS cyclases including the C-terminal domain of BenH from the benastatin BGC^27, 28^, which also comprises an N-terminal TcmN like cyclase domain (pfam 03364); and WhiE Protein 1 and other members of the SchA/CurD-like family of PKS enzymes commonly associated with BGCs for the biosynthesis of spore pigments in *Streptomyces* spp^29, 30, 31^. SchA/CurD- and WhiE Protein 1-like enzymes are comprised of an N-terminal ABM domain and a C-terminal PKS cyclase domain (pfam 00486; superfamily cl24023).

To interrogate their roles, we used Cas9-mediated genome editing^32^ to make in-frame deletions in each of these five putative cyclase genes (*forD, forL, forR, forS* and *forU*). Three independent mutants generated from each gene deletion experiment, along with the WT strain, were grown on MS agar and cultured at 30°C for nine days. To assess differences in secondary metabolite production the ethyl acetate extracts from each culture were subjected to HPLC(UV) and LCMS analysis (Supplementary Dataset 2). The metabolic profiles showed that the *ΔforD, ΔforL, ΔforR* and *ΔforU* mutants lost the ability to produce formicapyridines, fasamycins and formicamycins (Supplementary Figures 5-9 respectively), and no new shunt metabolites could be identified despite rigorous interrogation of the LCMS and LC(UV) data. These deletions could be rescued by complementation with the deleted gene under control of either the native promoter (*ΔforD/forD* and *ΔforL/forL*) or the constitutive *ermE*\* promoter (Δ*forR/forR*), although the titres did not reach that of the WT strain in all cases. For the Δ*forU* mutant complementation with *forU* restored production of the non-halogenated congener **10** only, indicating a polar effect on the downstream halogenase *forV* gene. Subsequent complementation with a *forUV* cassette under the *ermE*\* promoter in which the two genes were transcriptionally fused led to the production of halogenated fasamycin and formicamycin congeners.

In contrast, production of formicapyridines was increased approx. 25-fold in the Δ*forS* mutant when compared to the WT (Fig. 6). These effects were complemented by ectopic expression of *forS* under the control of the *ermE*\* promoter. Moreover, the Δ*forS* mutant was significantly compromised in its ability to produce formicamycins, with their titre being reduced to approximately one third that of the WT strain. This result is consistent with our hypothesis for formicapyridine biosynthesis and suggests that ForS plays a role during B-ring closure, and that this constitutes the final step of fasamycin backbone biosynthesis. However, it also demonstrates that the final cyclization step can occur reasonably efficiently without ForS. While carrying out analysis of the Δ*forS* mutant we also identified a new minor congener which was not otherwise identified in any WT strain fermentations. Scale up growth (4 L) and solvent extraction, followed by isolation (3.4 mg) and structural elucidation using the approaches described above, identified this compound (**13**) as the C24-carboxyl analogue of fasamycin C (which we have named fasamycin F). This is the first fasamycin congener identified with the C24-carboxyl group intact.

## Discussion

Using targeted metabolomics, we identified a new family of pyridine containing polyketide natural products that we have named the formicapyridines. Remarkably, these compounds are derived from the fasamycin/formicamycin biosynthetic machinery meaning that the *for* BGC is responsible for the production of three structurally differentiated pentacyclic scaffolds. Then, inspired by evolutionary considerations, we introduced a gene deletion into the BGC which significantly altered the relative levels of these metabolites in a targeted manner: the titre of formicapyridines was significantly increased (approx. 25-fold) in contrast to the fasamycin/formicamycins which were decreased to approx. one third of the WT titre. These results raise a series of questions about the cyclization events associated with the fasamycin/formicamycin biosynthetic pathway, as well as the maintenance of pathway fidelity.

Our data suggest that formation of ring-B of the fasamycin scaffold is the final biosynthetic step leading to the linear tetracyclic portion of the molecule, followed by thioester hydrolysis to liberate the ACP and subsequent decarboxylation to remove the carboxyl group attached to C24 (Fig. 5). Consistent with this, deletion of *forS* led to the isolation of a new shunt metabolite fasamycin F (**13**) with a C24-carboxyl group that was not observed from the WT. However, while ForS is implicated in ring-B cyclization it is not required for this role, nor, apparently, for the biosynthesis of any congeners. Rather, ForS seems to decrease the production of ‘aberrant’ congeners from the pathway, e.g. formicapyridines, while increasing overall productivity. The most parsimonious interpretation of these data is that another of the *for* BGC gene products is the actual catalyst for ring-B formation, and that ForS acts as a ‘chaperone’ which modulates or stabilises the assembly, or arrangement, of a multienzyme complex to optimise production of the fasamycins, and therefore ultimately the formicamycins. This has the consequence of minimising the production of shunt metabolites, i.e. the formicapyridines. Thus, while deletion of *forS* leads to the phenotype desired from our thought experiment, the mechanism by which this occurs is not what was anticipated. Interpretation of the *for* BGC bioinformatic analysis above suggests the most likely candidate for a ring-B (final) cyclase is ForL, due to its close relationship with TcmI which catalyses a similar final cyclization step during tetraceomycin biosynthesis^20, 21, 22^. The apparent lack of any intermediates or shunt metabolites being accumulated by the remaining mutants is somewhat surprising and suggests an absolute requirement for the formation of a PKS-cyclase complex before biosynthesis can be initiated.

**Figure 5.**
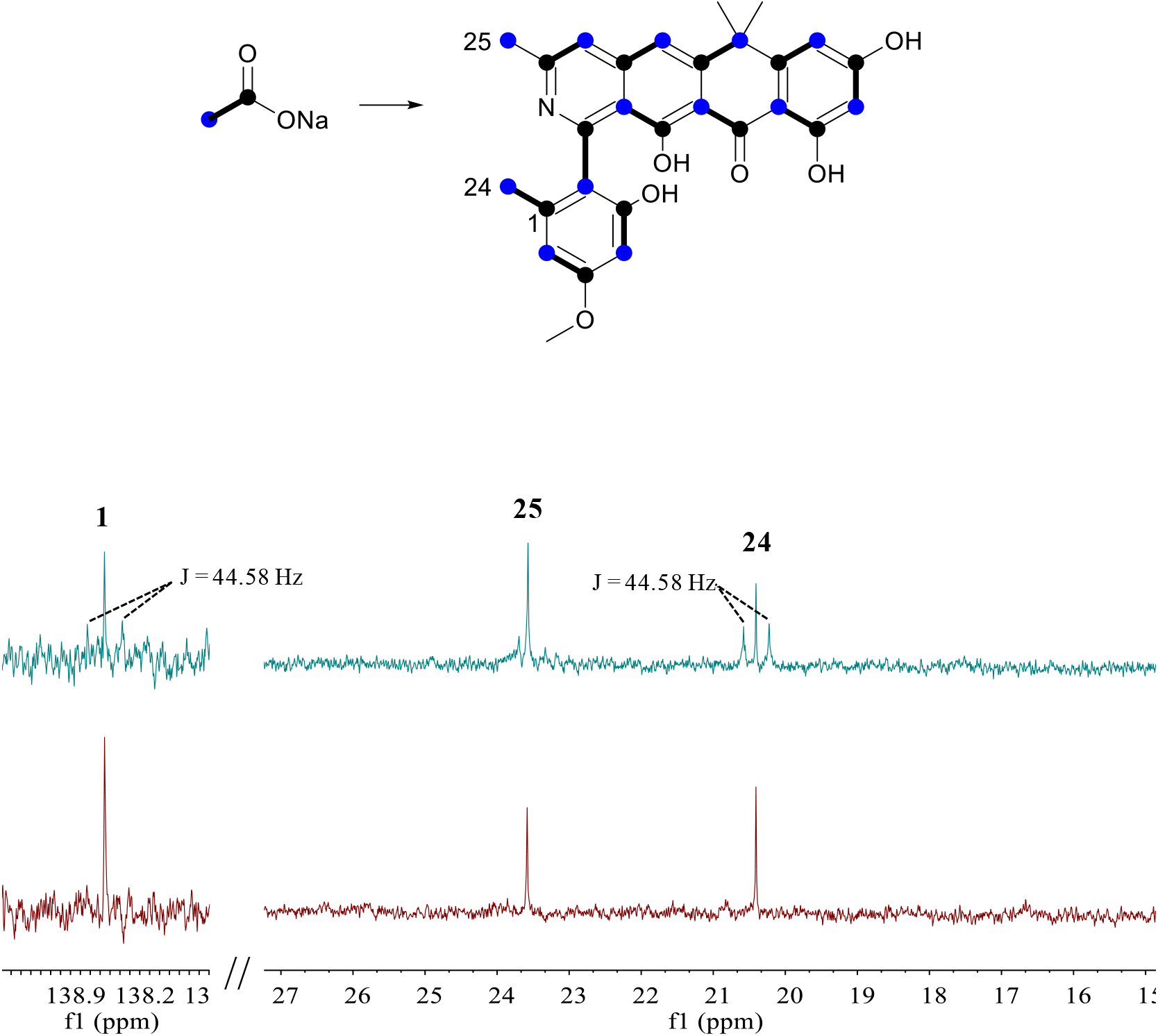
The methyl group carbon C25 arises from C2 of acetate. Comparison of ^13^C NMR spectra for **2** isolated after growth in the presence of [1,2-^13^C_2_] sodium acetate. Integration of the signal for C25 confirmed enrichment of the ^13^C isotope and the absence of coupling to any adjacent carbon atom. In contrast, the methyl group atom C24 shows enrichment and coupling to C1. The C2 atom of [1,2-^13^C_2_] sodium acetate is highlighted as a blue circle, the C1 atom as a black circle, and the coupled unit by a bold line.

**Figure 6.**
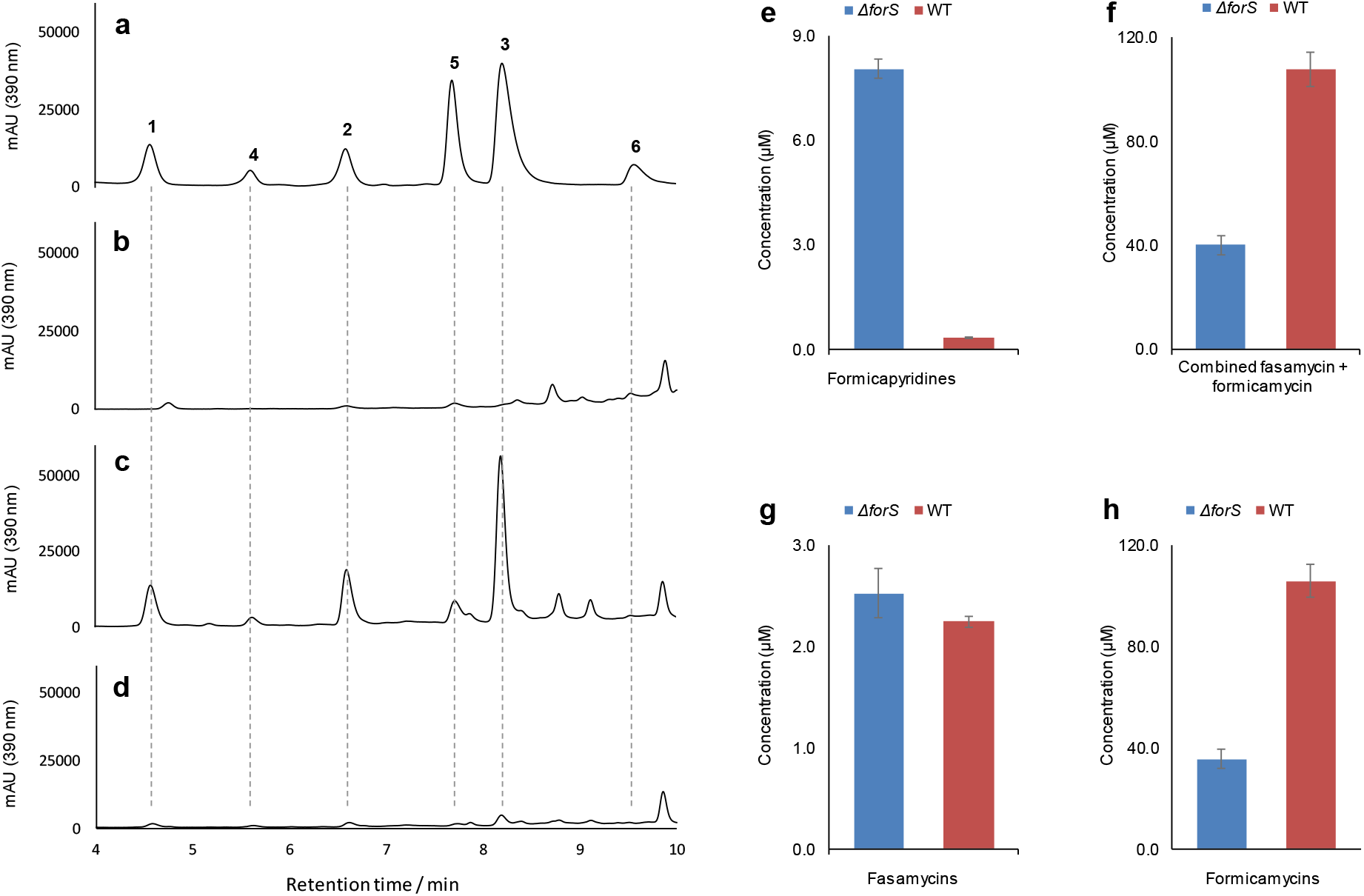
Mutational analysis of *forS*. Reconstituted HPLC chromatograms (UV; 390 nm) showing: (a) formicapyridine standards **1-6**; (b) *S. formicae* WT extract; (c) *S. formicae* Δ*forS* extract; (d) *S. formicae ΔforS/forS* extract. Quantitative data for the combined titre of each metabolite family produced by the *S. formicae* WT and Δ*forS* mutants are shown for: (e) total formicapyridines; (f) combined total fasamycins and formicamycins; (g) total fasamycins; (h) total formicamycins (mean ± standard deviation; n = 3).

These observations are reminiscent of studies regarding the biosynthesis of pradimicin^33^. The pradimicins are pentangular structures similar to the benastatins^27, 28^, and the BGC responsible for their production contains a complement of three PKS cyclases equivalent to ForD (PdmD), ForL (PdmK), and ForR (PdmL)^33, 34^ It also contains two ABM domain proteins (PdmH and PdmI; c.f. ForS and ForU) which were assigned monooxygenase roles. Through heterologous expression of PKS gene cassettes it was deduced that for the biosynthetic pathway to function correctly, and yield a pentangular backbone, all three cyclase genes (PdmDKL) plus the ABM domain monooxygenase PdmH must be co-expressed. It should be noted that in this pathway an oxidation reaction to form the quinone structure is required. These results led the authors to propose a model in which the two cyclases PdmKL and the monooxygenase PdmH form a multienzyme complex that engulfs the entire polyketide molecule during its assembly and work synergistically to ensure the correct reaction pathway occurs thereby minimising the production of shunt metabolites^33^. Similarly, formation of the unusual discoid metabolite resistomycin involves an extremely rare S-shaped folding pattern and requires the coordinated function, likely as a multienzyme complex, of core PKS proteins in addition to three distinct cyclase enzymes^24^. Moreover, heterologous reconstruction of the resistomycin pathway gave no products when the minimal PKS plus first cyclase were assembled^24^, a rare observation that is in keeping with our data showing the requirement of all the putative cyclases ForDLRU to produce any pathway derived metabolites, including shunts.

During biosynthesis of the *for* BCG polyketide backbones there is no requirement for the function of a monooxygenase. Consistent with this, deletion of *forS* does not abolish polyketide production, but instead affects pathway productivity and fidelity. Thus, based on our mutational data, and the observations discussed above, we hypothesise that ABM family enzymes can act as monooxygenases and/or as ancillary proteins to tune, in some way, the PKS enzyme complex function, and therefore the biosynthetic pathway, during aromatic polyketide biosynthesis. Genes encoding these proteins occur commonly in PKS BGCs, and multiple paralogues are often present, even when monooxygenase reactions are not required. It is noteworthy that several PKS cyclases occur as fusion proteins with an ABM domain, which can be located at either terminus. We speculate, tentatively, that these may represent examples of mature pathways where the chaperone-like function of the ABM family protein has become essential, leading to selective pressure for the encoding gene to become transcriptionally fused with other cyclase encoding genes.

## Materials & Methods

### Chemistry methods and materials

Unless stated otherwise all chemicals were supplied by Sigma-Aldrich or Fisher Scientific. [1,2-^13^C_2_] sodium acetate was purchased from CORTECNET. All solvents were of HPLC grade or equivalent. NMR spectra were recorded on a Bruker Avance III 400 MHz NMR spectrometer equipped with 5 mm BBFO Plus probe unless noted otherwise. The ^13^C NMR spectra for **2** and isotopically enriched **2** after feeding [1,2-^13^C_2_] sodium acetate were recorded on a Bruker Avance III 500 MHz NMR spectrometer equipped with a DUL cryoprobe at 30°C.

Unless otherwise stated samples were analysed by LCMS/MS on a Nexera/Prominence UHPLC system attached to a Shimadzu ion-trap time-of-flight (IT-ToF) mass spectrometer. The spray chamber conditions were: heat-block, 300°C; 250° curved desorbtion line; interface (probe) voltage: 4.5 KV nebulizer gas flow rate 1.5 L/min; drying gas on. The instrument was calibrated using sodium trifluoroacetate cluster ions according to the manufacturer’s instructions and run with positive-negative mode switching. The following analytical LCMS method was used throughout this study unless otherwise stated: Phenomenex Kinetex C18 column (100 × 2.1 mm, 100 Å); mobile phase A: water + 0.1% formic acid; mobile phase B: methanol. Elution gradient: 0–1 min, 20% B; 1–12 min, 20%–100% B; 12–14 min, 100% B; 14–14.1 min, 100%–20% B; 14.1–17 min, 20% B; flow rate 0.6 mL/min; injection volume 10 μL. Samples for were prepared for LCMS analysis by taking a rectangle of agar (2 cm^3^) from an agar plate culture and shaking with ethyl acetate (1 mL) for 20 min. The ethyl acetate was transferred to a clean tube and the solvent removed under reduced pressure. The resulting extract was dissolved in methanol (200 μL).

### Standard microbiology and molecular biology methods

All strains used or made in this study are described in Supplementary Table 1. All plasmids and ePACs used are described in Supplementary Table 2. All PCR primers used are described in Supplementary Table 3. The composition of media used are described in Supplementary Table 4. The antibiotics and their concentrations used are described in Supplementary Table 5. Standard DNA sequencing was carried out by Eurofins Genomics using the Mix2Seq kit (Ebersberg, Germany).

*E. coli* strains were cultivated at 37°C in LB Lennox Broth (LB), shaking at 220 rpm, or LB agar supplemented with antibiotics as appropriate. *S. formicae* was cultivated at 30°C on mannitol soya flour (MS) agar or MYM agar with appropriate antibiotic selection. To prepare *Streptomyces* spores, material from a single colony was plated out using a sterile cotton bud and incubated at 30°C for 7-10 days until a confluent lawn had grown over the entire surface of the agar. Spores were harvested by applying 20% glycerol (2 mL) to the surface of the agar plate culture and gently removing spores with a sterile cotton bud before storing them at −80°C. Glycerol stocks of *E. coli* were made by pelleting the cells from an overnight *E. coli* culture (3-5 mL) in a bench top centrifuge (5000 rpm, 5 min) and resuspending in fresh, sterile 1:1 2YT/40% glycerol (1 mL). Glycerol stocks were stored at −80°C.

*S. formicae* genomic DNA and cosmid ePAC DNA (from *E. coli* DH10B) was isolated using a phenol:chloroform extraction method. Briefly, cells from an overnight culture (1 mL) were pelleted at 13,000 rpm in a benchtop microcentrifuge and resuspended in solution 1 (100 μL) (50 mM Tris/HCl, pH 8; 10 mM EDTA). Alkaline lysis was performed by adding solution 2 (200 μL) (200 mM NaOH; 1%SDS) and mixing by inverting. Solution 3 (150 μl) (3M potassium acetate, pH 5.5) was added and samples mixed by inverting, before the soluble material was harvested by microcentrifugation at 13,000 rpm for 5 min. The nucleic acid was extracted with 25:24:1 phenol:chloroform:isoamyl alcohol (400 μL), and DNA was precipitated in 600 μl ice-cold isopropanol. After centrifugation, the resulting DNA pellet was washed in 200 μl 70% ethanol and air dried before being resuspended in water for quantification on a Nanodrop 2000c UV-Vis spectrophotometer. Plasmid DNA was isolated from *E. coli* strains using a Qiagen miniprep kit according to the manufacturer’s instructions. PAC pESAC13-215-G DNA, containing the entire *for* BGC, was used as the template for all PCR reactions. DNA amplification for cloning was conducted using Q5 Polymerase and diagnostic PCR was set up using PCRBIO Taw Mix Red (PCR Biosystems), as per the manufacturers’ instructions. Amplified fragments and digested DNA products were purified on 1% agarose gels by electrophoresis and extraction using the Qiagen gel extraction kit as per the manufacturer’s instructions. For overlay bioassays soft LB agar (100 mL LB with 0.5% agar) was inoculated with *Bacillus subtillis* 168 (10 mL; approx. OD_600_ = 0.6). Set concentrations of the compound for testing were made up in methanol and an aliquot (20 μL) was applied to Whatman 6mm antibiotic assay discs, air dried, and the discs placed in the centre of the solidified agar plates. Plates were incubated at 30°C overnight before being examined for zones of inhibition around the discs. For liquid culture bioassays, cultures of *B. subtillis* 168 (1 mL) were grown at 30°C, 200 rpm shaking, with the relevant antibiotic (50 μg/mL). After 7 hours incubation, samples were taken from the cultures, diluted in series and plates for colony count in triplicate following Miles and Misra protocol^35^.

### Generating mutant strains of *S. formicae*

CRISPR/Cas9 genome editing was conducted as described previously using the pCRISPomyces-2 plasmid supplied by Addgene^1,2^. Protospacers to use in the synthetic guide RNA (sgRNA) were annealed by heating to 95°C for 5 min followed by ramping to 4°C at 0.1°C/sec. Annealed protospacers were assembled into the pCRISPomyces-2 backbone at the BbsI site by Golden Gate assembly as described previously^5^. The two homology repair template arms (each approx. 1kb) were assembled into the plasmid containing the sgRNA at the *XbaI* site using Gibson Assembly as described previously^5^. Genetic complementation was achieved using either the native promoter or the constitutive, high-level *ermE*\* promoter and single copies of the relevant gene(s) cloned into the integrative vector pMS82, as described previously^5^. Gibson assembly was used to fuse the gene(s) (and the native promoter if located distally in the BGC) and assemble them into the chosen plasmid. Plasmids were confirmed by PCR amplification and sequencing. Plasmids were then conjugated into *S. formicae* KY5 via the non-methylating *E. coli* strain ET12567 containing pUZ8002, as described previously^5, 36^. Ex-conjugants were selected on the appropriate antibiotics and plasmids were cured from *S. formicae* using temperature selection at 37°C.

### Production, purification and structure determination of formicapyridines

*S. formicae* was cultivated on MS agar (14 L; approx. 450 plates) at 30°C for nine days. The agar was sliced into small pieces and extracted twice with ethyl acetate (10 L) using ultrasonication to improve the extraction. The extracts were combined, and the solvent removed under reduced pressure to yield a brown oil which was dissolved in methanol (20 mL). This extract was first chromatographed over a Phenomenex Gemini-NX reversed-phase column (C18, 110 Å, 150 × 21.2 mm) using a Thermo Scientific Dionex Ultimate 3000 HPLC system and eluting with the following gradient method: (mobile phase A: water + 0.1% formic acid; mobile phase B: acetonitrile) 0–5 min 40% B; 5–35 min 40%-100% B; 35–40 min 100% B; 40–40.1 min 100%-40% B; and 40.1–45 min 40% B; flowrate 20 mL/min; injection volume 1 mL. Absorbance was monitored at 250 nm. Fractions (20 mL) were collected and analysed by LCMS. Fractions 2 to 4 contained **1-6** and were further purified by chromatography over a Phenomenex Gemini-NX semi-prep reversed-phase column (C_18_, 110 Å, 150 × 10 mm) using an Agilent 1100 series HPLC system and eluting with the following gradient method: (mobile phase A: water + 0.1% formic acid; mobile phase B: acetonitrile) 0–2 min 40% B; 2–20 min 40%–100% B; 20–21 min 100% B; 21–21.1 min 100%–40% B; 21.1–23 min 40% B; flowrate 3 mL/min; injection volume 100 μL). Absorbance was monitored at 390 nm. The samples were finally purified by Sephadex LH20 size exclusion chromatography with 100% methanol as the mobile phase. The isolated yields were: **1** (1 mg), **2** (2 mg), **3** (2 mg), **4** (0.7 mg), **5** (1 mg) and **6** (0.6 mg). These pure compounds were subjected to analysis by HRMS and 1D and 2D NMR as described in the main text (see Figs 1 and 2). Spectroscopic and other data for each compound is presented in Supplementary Note 2.

### Stable isotope feeding experiment

*S. formicae* was cultivated on MS agar (3 L; approx. 100 plates) at 30°C and overlaid with [1,2-^13^C_2_] sodium acetate (1 mL of a 60 mM solution) after 24 h, 48 h, 72 h, 96 h and 120 h. After a further 72 h the agar was extracted and purified using the methods described above to yield a sample of **2** (0.9 mg). This material was analyzed by LCMS and ^13^C NMR (125 MHz; 4096 scans; d4-methanol). However, due to the weak and overlapping signals, only the following coupling constants (*J_CC_*) of the intact acetate units were recorded: C24-C1, 44.61 Hz; C2-C3, 68.58 Hz; C4-C5, 68.16Hz; C20-C21, 56.18 Hz. In addition, C14, C16, C18, and C22 have coupling constant of 66.47, 67.61, 42.08 and 60.43 Hz respectively. Spectroscopic data is presented in Fig. 5 and Supplementary Figure 4.

### Production, purification and structure determination of fasamycin F (13)

*S. formicae ΔforS* was cultivated on 4L (~120 plates) of MS agar at 30°C for nine days. The agar was extracted and purified using the methods described above to yield **13** (3.4 mg). This material was analysed by LCMS and ^13^C NMR (100 MHz; 6500 scans; *d*_4_-methanol). Spectroscopic data is presented in Supplementary Note 2.

### Chemical analysis of congener content in cyclase mutants

*S. formicae* wild type or mutant strains (n=3) were grown on MS agar at 30°C for nine days. A rectangle of agar (2 cm^3^) was excised from each petri dish, sliced into small pieces and shaken with ethyl acetate (1 mL) for 20 min. The ethyl acetate was transferred to a clean tube and the solvent removed under reduced pressure. The resulting extract was dissolved in methanol (200 μL) and analysed by LCMS but using the following modified UPLC method: Phenomenex Gemini C18 column (100 × 2.1 mm, 100 Å); mobile phase A: water + 0.1% formic acid; mobile phase B: methanol. Elution gradient: 0–2 min, 50% B; 2–14 min, 50%–100% B; 14–18 min, 100% B; 18–18.1 min, 100%–50% B; 18.1–20 min, 50% B; flow rate 1 mL/min; injection volume 10 μL.

Calibration curves (Supplementary Figure 10; Supplementary Dataset 2) were determined using standard solutions of fasamycin C **10** (10, 50, and 200 μM), formicamycin C **16** (10, 50, and 200 μM), formicapyridine D **4** (5, 10, 25, and 50 μM) and fasamycin F **13** (5, 10, and 100 μM). The content of **10** and **16** was determined by UV absorption at 285 nm. The content of **4** and **13** was determined by MS analysis of the base peak chromatogram (positive mode).

## Data availability

The authors declare that the data supporting the findings reported in this study are available within the article and the Supplementary Information or are available from the authors on reasonable request.

## Supporting information

Supplementary data set 1

Supplementary data set 2

## Acknowledgements

This work was supported by the Biotechnology and Biological Sciences Research Council (BBSRC) *via* Institute Strategic Program Project BBS/E/J/000PR9790 to the John Innes Centre (ZQ) and by a Norwich Research Park BBSRC Postdoctoral Training Program Studentship BB/M011216/1 (RD). We thank Dr Lionel Hill and Dr Gerhard Saalbach (JIC) for excellent metabolomics support. We dedicate this manuscript to our deceased colleague Dr Karl A. Wilkinson (University of Cape Town) who provided helpful input and enlightening discussions during the early phases of this work.

## Author contributions

Z.Q., R.D., B.W. and M.I.H. designed the research. Z.Q., R.D., B.W. and M.I.H. wrote the manuscript and all authors commented. Z.Q. performed the chemical experiments, R.D. performed the molecular genetics experiments.

## Competing interests

The authors declare no competing financial interests.

