## Supplementary information for "Aromatic polyketide biosynthesis: fidelity, evolution and engineering"

### Electronic Supporting information

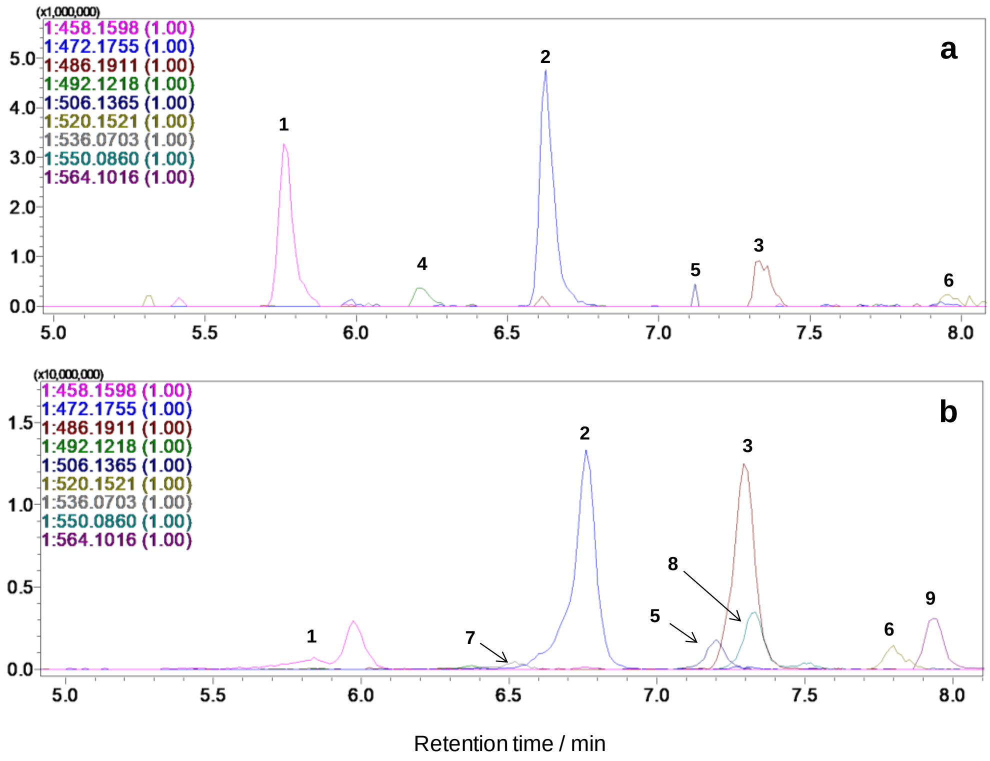


**Supplementary Figure 1 |** Extracted ion chromatograms for **1**-**9** from LCMS runs of extracts from the WT strain grown on MS agar medium without (a) and with (b) exogenous addition of NaBr (the slight shifts in retention times for **1**-**6** was due to machine maintenance carried out between the experiments).

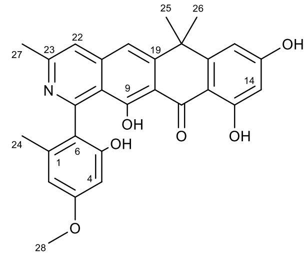


Formicapyridine B (**2**)

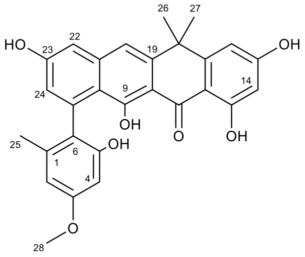


Fasamycin C (**10**)

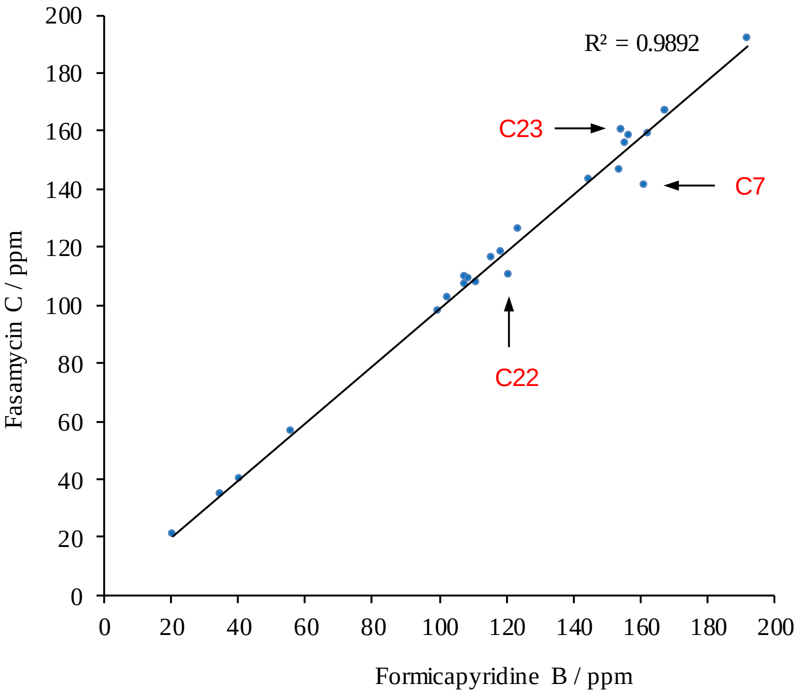


**Supplementary Figure 2 |** Comparison of the $^{13}C$ NMR chemical shifts for formicapyridine B (**2**) and fasamycin C (**10**). The chemical shifts for carbons C7, C22 and C23 are highlighted.

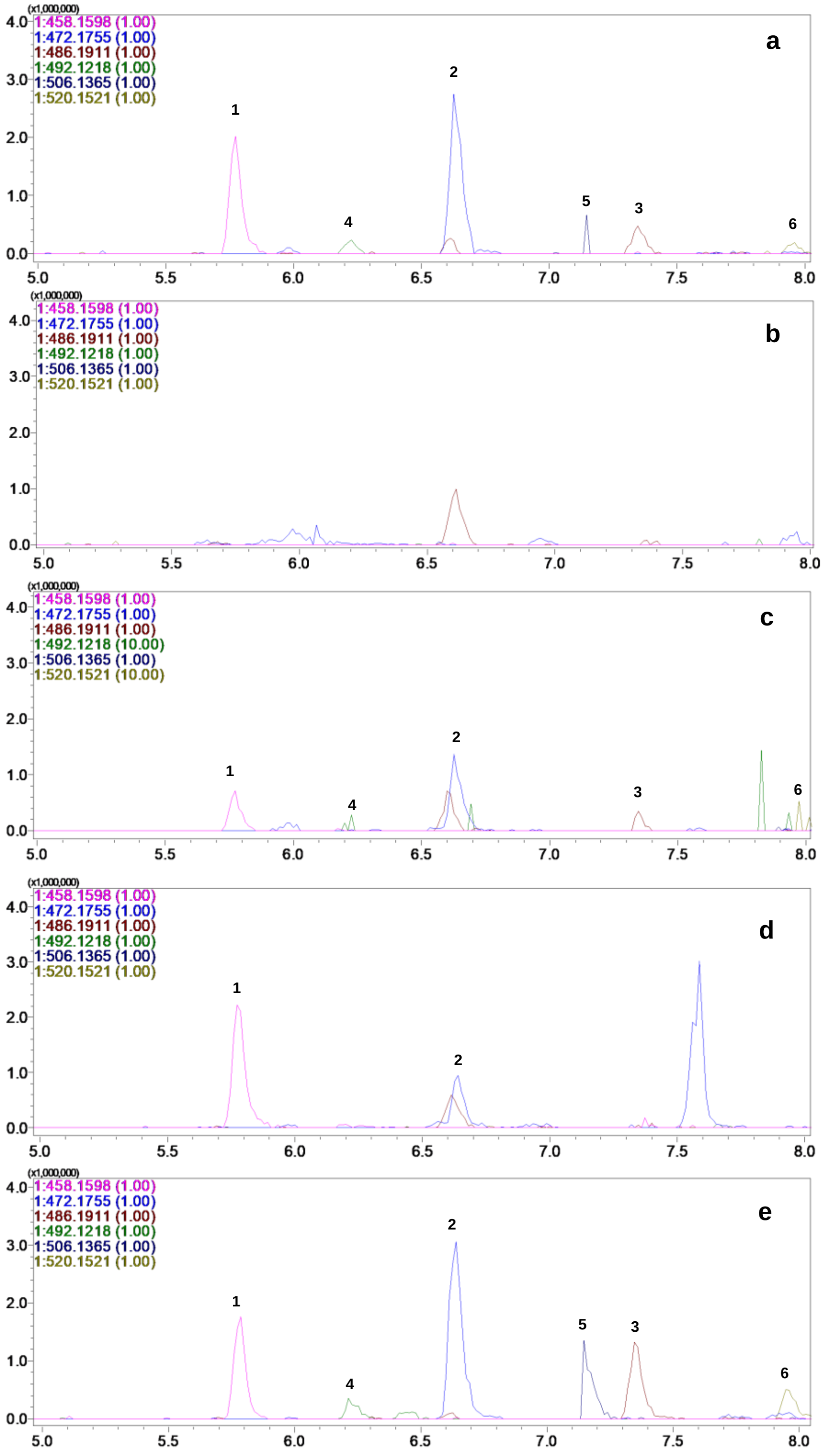


**Supplementary Figure 3 |** Extracted ion chromatograms for **1**-**6** from LCMS runs of fermentation extracts for: (a) *S. formicae* WT; (b) *S. formicae* Δ*for*; (c) *S. formicae* Δ*for/for*; (d) *S. formicae* Δ*forV*; (e) *S. formicae* Δ*forv/forv*. (The signal abundance of **4** and **6** in (c) was enlarged × 10 for clarity).

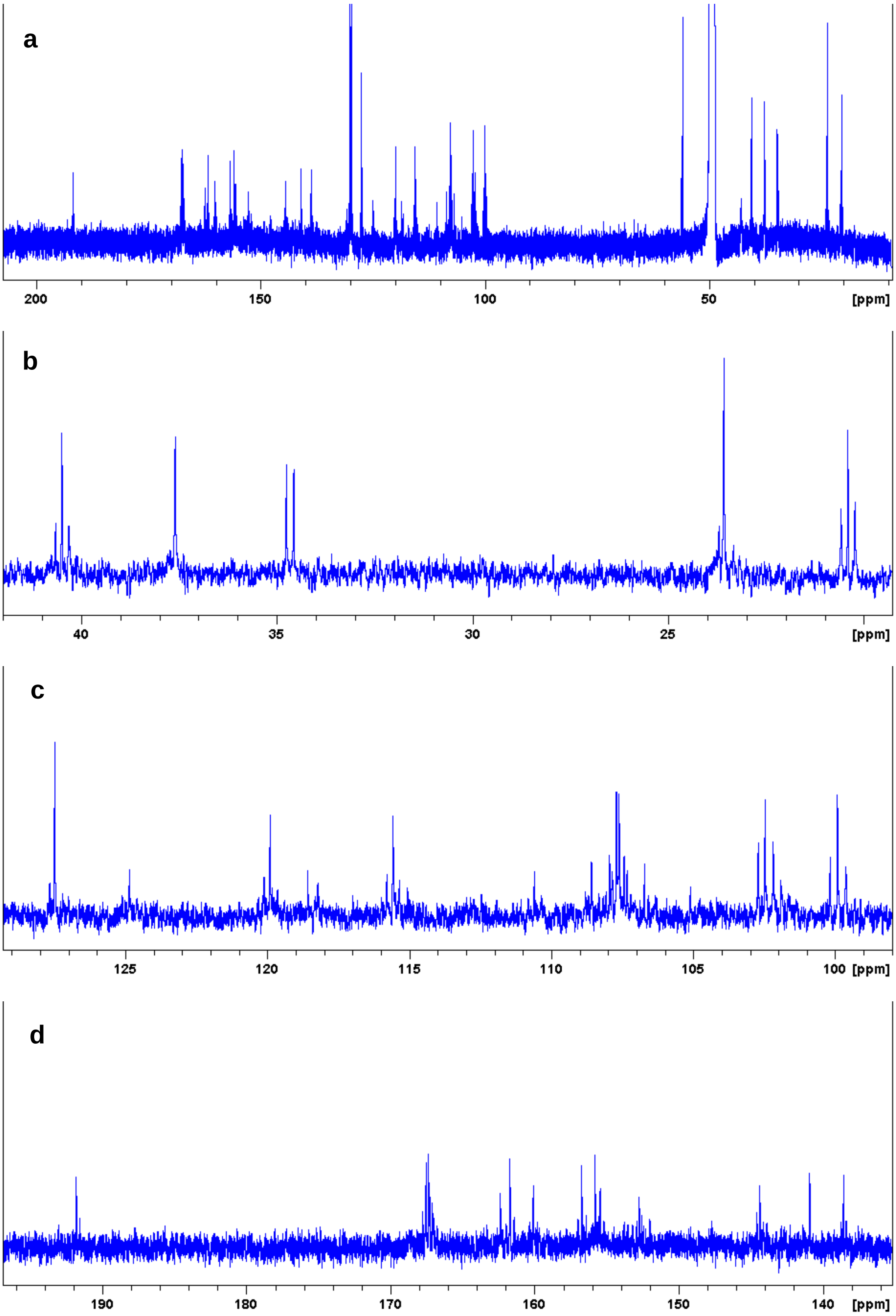


**Supplementary Figure 4 |** $^{13}C$ NMR spectra (125 MHz, methanol-$d_6$) of **2** isolated after growth of *S. formicae* WT in the presence of [1,2-$^{13}C_2$] sodium acetate: (a), full scale; (b)-(d), expanded scale.

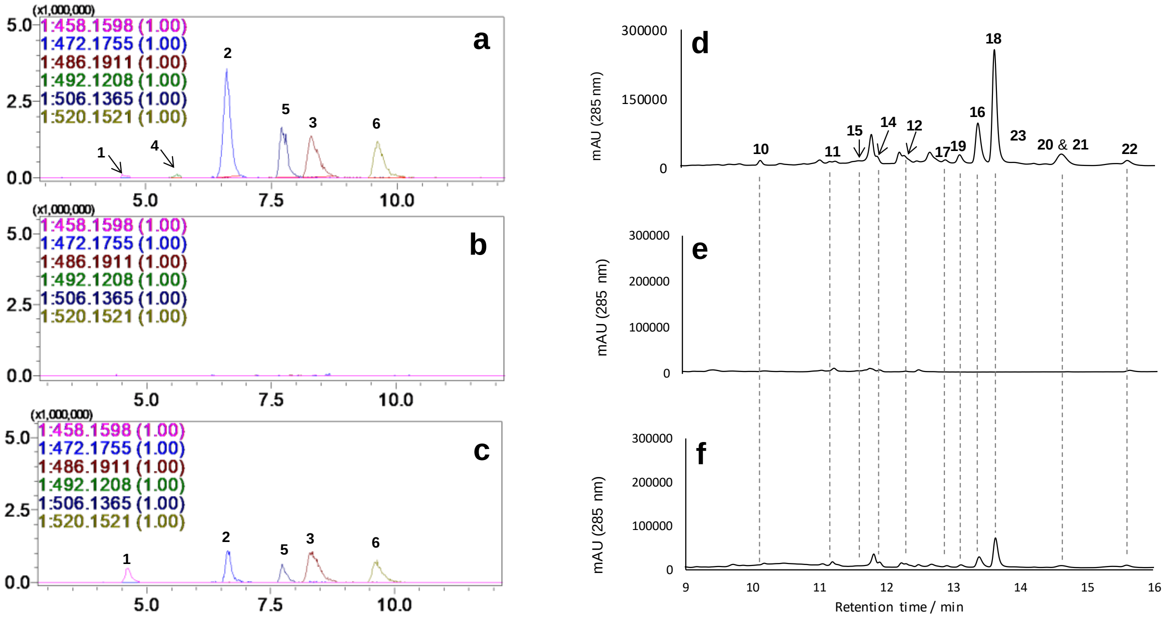


**Supplementary Figure 5 | Mutational analysis of *forD*.** Extracted ion chromatogram (left) and reconstituted HPLC-UV (285 nm) profiles (right) for extracts of *forD* mutagenesis and complementation experiments: (a) and (d) *S. formicae* wild-type; (b) and (e) *S. formicae* Δ*forD*; (c) and (f) *S. formicae* Δ*forD/forD*.

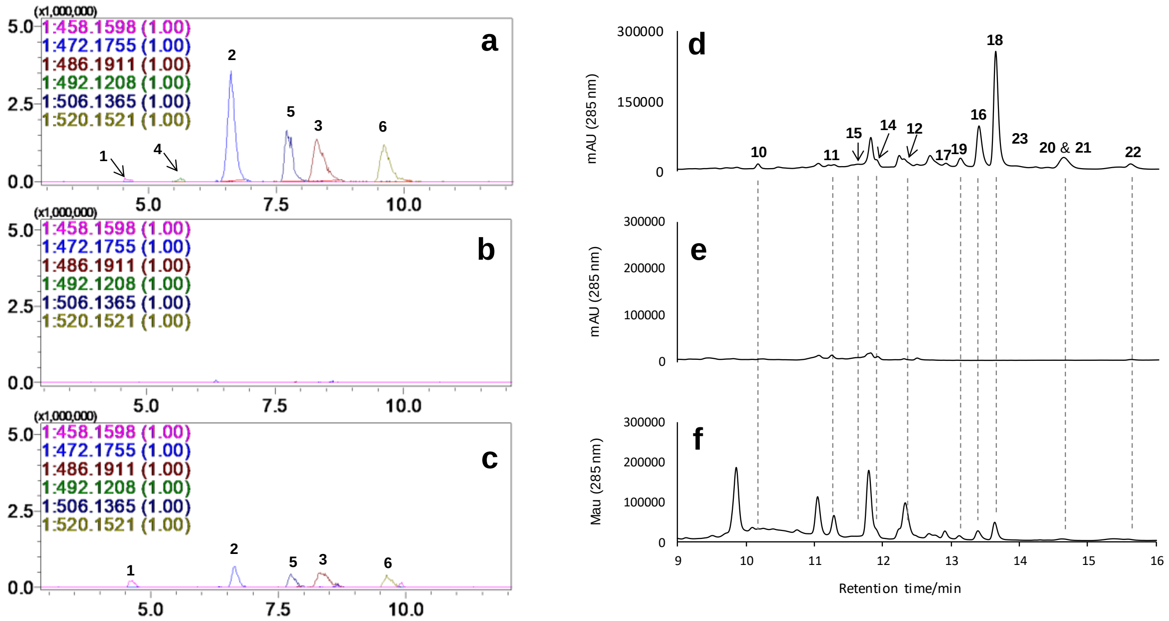


**Supplementary Figure 6 | Mutational analysis of *forL*.** Extracted ion chromatogram (left) and reconstituted HPLC-UV (285 nm) profiles (right) for extracts of *forL* mutagenesis and complementation experiments: (a) and (d) *S. formicae* wild-type; (b) and (e) *S. formicae* Δ*forL*; (c) and (f) *S. formicae* Δ*forL/forL*.

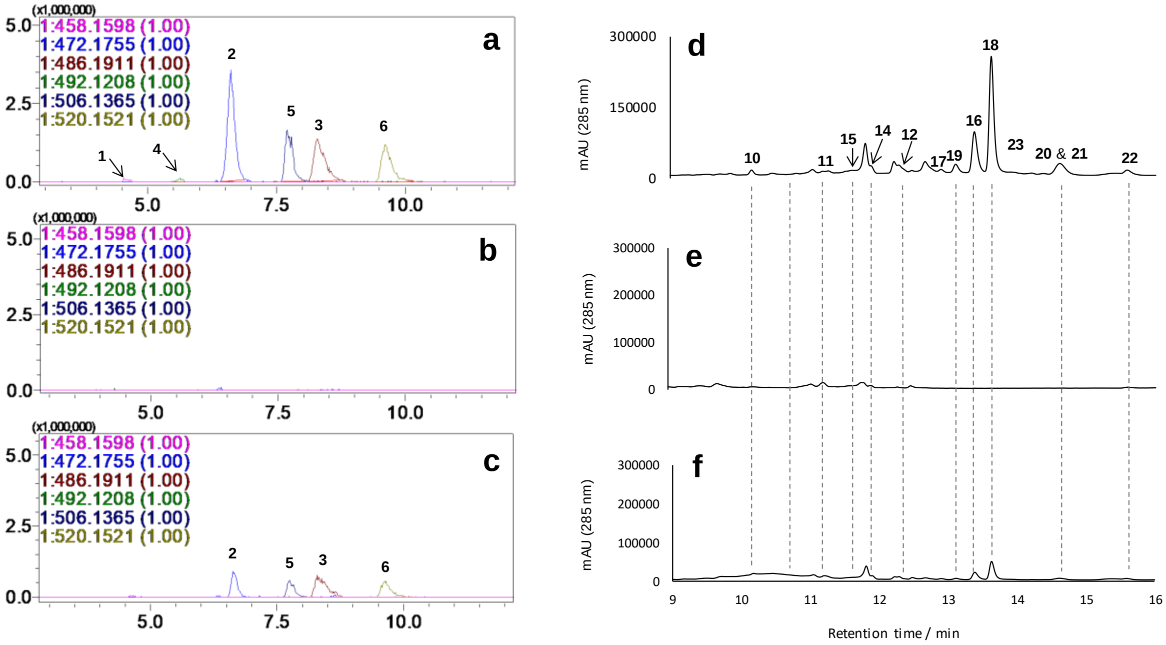


**Supplementary Figure 7 | Mutational analysis of *forR*.** Extracted ion chromatogram (left) and reconstituted HPLC-UV (285 nm) profiles (right) for extracts of *forR* mutagenesis and complementation experiments: (a) and (d) *S. formicae* wild-type; (b) and (e) *S. formicae* Δ*forR*; (c) and (f) *S. formicae* Δ*forR/forR*.

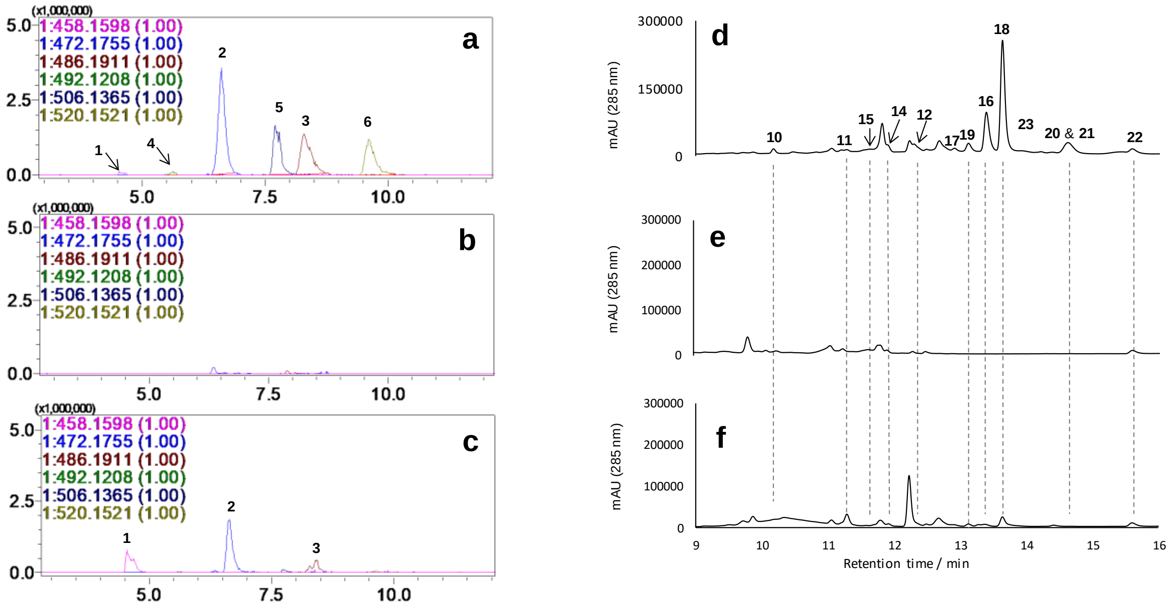


**Supplementary Figure 8 | Mutational analysis of *forU*.** Extracted ion chromatogram (left) and reconstituted HPLC-UV (285 nm) profiles (right) for extracts of *forU* mutagenesis and complementation experiments: (a) and (d) *S. formicae* wild-type; (b) and (e) *S. formicae* Δ*forU*; (c) and (f) *S. formicae* Δ*forU/forUV*.

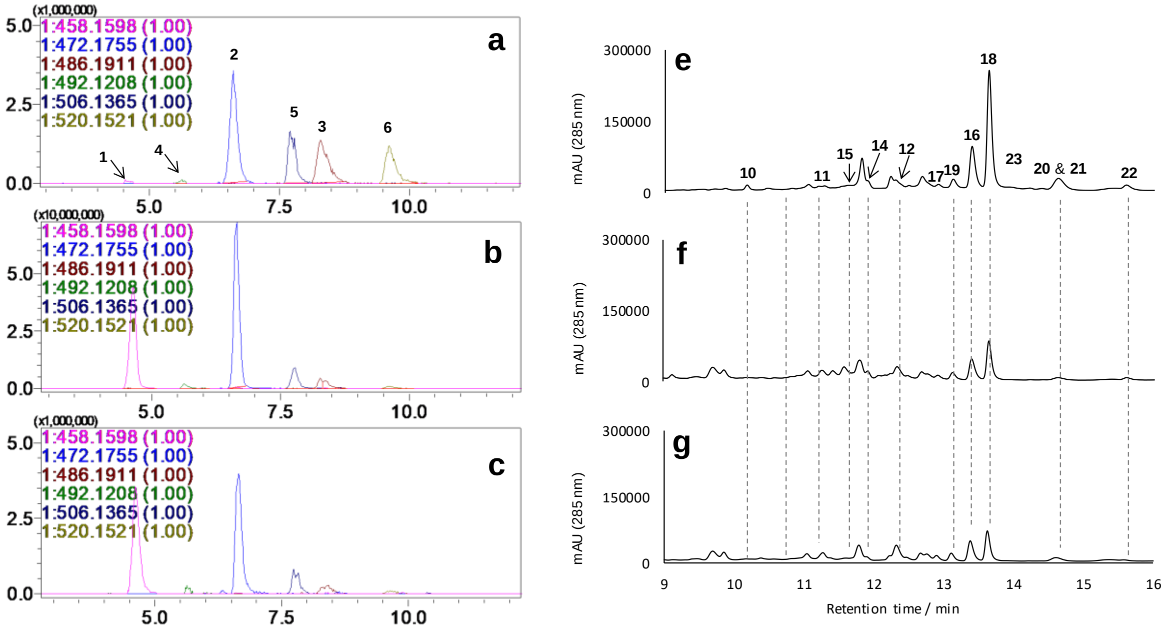


**Supplementary Figure 9 | Mutational analysis of *forS*.** Extracted ion chromatogram (left) and reconstituted HPLC-UV (285 nm) profiles (right) for extracts of *forS* mutagenesis and complementation experiments: (a) and (d) *S. formicae* wild-type; (b) and (e) *S. formicae* Δ*forS*; (c) and (f) *S. formicae* Δ*forS/forS*.

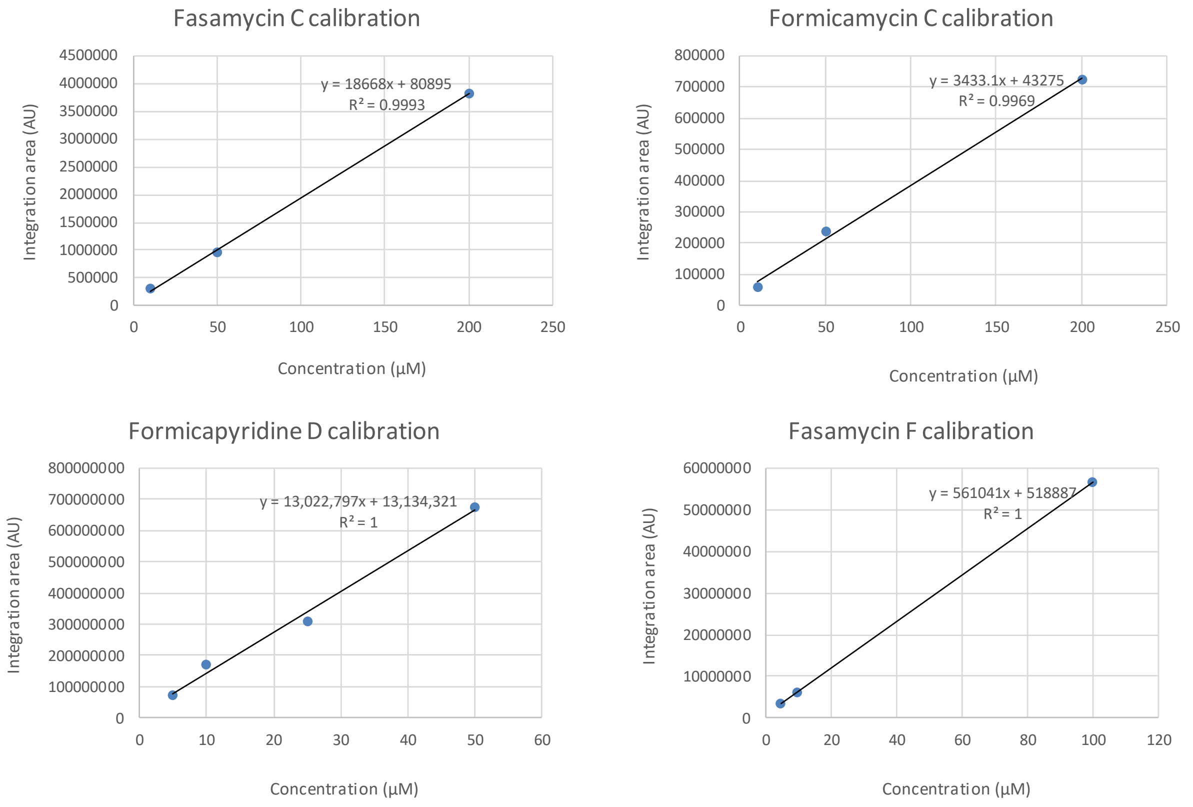


**Supplementary Figure 10 | Calibration curves of fasamycin C (10), formicamycin C (16), formicapyridine D (4), and fasamycin F (13).**

**Supplementary Table 1 | Strains made or used in this study.**

| Strain | Description | Plasmid | Resistance | Source or Reference |
|---|---|---|---|---|
| *E. coli* ET12567 | *dam*⁻ *dcm*⁻ *hsdS*⁻ | *pUZ8002* | *Cml*$^R$/*Tet*$^R$ | Reference 1 |
| *E. coli* Top10 | F– *mcr*A Δ(*mrr-hsd*RMS*mcr*BC) Φ80*lac*ZΔM15 Δ*lac*X74 *rec*A1 *end*A1 *ara*D139 Δ(*ara leu*) 7697 *gal*U *gal*K *rps*L *nup*G | | | Invitrogen, USA |
| *E. coli* DH10ß | F– *mcr*A Δ(*mrr-hsd*RMS*mcr*BC) Φ80*lac*ZΔM15 Δ*lac*X74 *rec*A1 *end*A1 *ara*D139 Δ(*ara leu*) 7697 *gal*U *gal*K *rps*L *nup*G λ– | | | Invitrogen, USA |
| *S. formicae* | Wild-type strain | | | Reference **Error! Bookmark not defined.** |
| *S. formicae* Δ*for* | Formicamycin (*for*) biosynthetic gene cluster (BCG) deleted | | | Reference **Error! Bookmark not defined.** |
| *S. formicae Δfor:for* ΦC31 | *for* BGC deleted and complemented with the *for* BGC in trans at ΦC31 | *PESAC-13 215-G* | *Kan*$^R$/*Tsr* | Reference **Error! Bookmark not defined.** |
| *S. formicae* Δ*forV* | Halogenase (*forV*) gene deleted | | | Reference **Error! Bookmark not defined.** |
| S. | Halogenase (*forV*) | pRD004 | Hyg$^R$ | Reference |

| | | | | |
|---|---|---|---|---|
| *formicae ΔforV:forV pForV* | deleted and complemented at ΦBT1 under the native promoter | | | **Error! Bookmark not defined.** |
| *S. formicae ΔforD* | Cyclase (*forD*) gene deleted | | | This work |
| S. *formicae ΔforD:forD pForD* | Cyclase (*forD*) deleted and complemented at ΦBT1 under the native promoter | pRD012 | $Hyg^R$ | This work |
| *S. formicae ΔforL* | Cyclase (*forL*) gene deleted | | | This work |
| S. *formicae ΔforL:forL pForL* | Cyclase (*forL*) deleted and complemented at ΦBT1 under the native promoter | pRD013 | $Hyg^R$ | This work |
| *S. formicae ΔforS* | Cyclase (*forR*) gene deleted | | | This work |
| S. *formicae ΔforS:forS pErmE** | Cyclase (*forR*) deleted and complemented at ΦBT1 under the ErmE* promoter | pRD015 | $Hyg^R$ | This work |
| *S. formicae ΔforR* | Cyclase (*forS*) gene deleted | | | This work |
| S. *formicae ΔforR:forR-A pErmE** | Cyclase (*forS*) deleted and complemented with *forR* and *forA* (directly downstream) at ΦBT1 under the ErmE* promoter | pRD017 | $Hyg^R$ | This work |
| *S. formicae ΔforU* | Cyclase (*forU*) gene deleted | | | This work |

| | | | | |
|---|---|---|---|---|
| S. *formicae* Δ*forU:forU-V pErmE** | Cyclase (*forU*) deleted and complemented with *forU* and *forV* (directly downstream) at ΦBT1 under the ErmE* promoter | pRD019 | $Hyg^R$ | This work |

**Supplementary Table 2 | Plasmids and ePACs used in this study.**

| **Plasmids and ePACs** | **Description** | **Source or Reference** |
|---|---|---|
| pCRISPomyces-2 | Streptomyces plasmid for expression of codon-optimized Cas9 and a single guide RNA | Reference 2 |
| pUZ8002 | Non-transmissible RK2 derivative with a mutation in *oriT* | Reference 3 |
| pESAC13-215-G | ePAC clone harbouring the *For* BGC integrative ΦC31 | Reference**Error! Bookmark not defined.**1 |
| pMS82 | ΦBT1 *attP-int* derived integration vector for the conjugal transfer of DNA from *E. coli* to *Streptomyces* ($Hyg^R$) | Reference 4 |
| pIJ10257 | Plasmid for the conjugal transfer of DNA (under control of the ermE* constitutive promoter) from E. coli to Streptomyces spp. Integrates specifically at the ΦBT1 attachment site (HygR) | Reference 5 |
| pRD004 | PMS82 pForV ForV complementation plasmid | This work |
| pRD012 | PMS82 pForD ForD | This work |

| | | |
|---|---|---|
| | complementation plasmid | |
| pRD013 | PMS82 pForL ForL complementation plasmid | This work |
| pRD015 | pIJ10257 ForR-ForA complementation plasmid | This work |
| pRD017 | pIJ10257 ForS complementation plasmid | This work |
| pRD019 | pIJ10257 ForU-ForV complementation plasmid | This work |

**Supplementary Table 3 | PCR primers used in this study.**

| **Primer name** | **Sequence** | **Description** |
|---|---|---|
| pCRISP-2 For TEST | AGGCTAGTCCGTTATCAACTTGAAA | Detection of pCRISP and cloning inserts around XbaI site forward |
| pCRISP-2 Rev TEST | TCGCCACCTCTGACTTGAGCGTCGA | Detection of pCRISP and cloning inserts around XbaI site reverse |
| Spacer primer | atacggctgccagataaggc | Detection of gRNA insert at BbsI site via sequencing |
| pms82 Rev | gccagtggtatttatgtcaacaccgcc | Detection of Pms82 and cloning inserts around HINDIII site reverse |
| FOR pMS82 | gcaacagtgccgttgatcgtgctatg | Detection of Pms82 and cloning inserts around HINDIII site forward |
| RD206 ForD 1.1 For | gctcggttgccgccgggcgtttttttaTCTAGAccagttcggccacggactgc | pCRISPomyces-2 template 1 left flank ForD |
| RD207 ForD 1.2 Rev | GCTGCTGCGACCAGGCGAGCTCGCcacggtcaggctcccttcg | pCRISPomyces-2 template 1 right flank ForD |
| RD208 ForD 2.1 For | GCGAGCTCGCCTGGTCGCAGCAGCgtatgagcgccaccgaggcc | pCRISPomyces-2 template 2 left flank |

| | | |
|---|---|---|
| | | ForD |
| RD209 ForD 2.2 Rev | gcaacgcggcctttttacggttcctggccTCTAGAccaacggccagacggcgcgtc | pCRISPomyces-2 template 2 right flank ForD |
| RD209 ForD gRNA For | ACGCtggatgcgcatgaacgcgaa | ForD gRNA forward |
| RD210 ForD gRNA Rev | AAACttcgcgttcatgcgcatcca | ForD gRNA reverse |
| RD211 ForD Test Int For | ccacgctggtcgaacagtgc | Confirmation of ForD deletion external for |
| RD212 ForD Test Int Rev | gcgagtctgaccaggcgctc | Confirmation of ForD deletion external rev |
| RD213 ForD Test Ext For | cgtcctggtcaccgacgagg | Confirmation of ForD deletion internal for |
| RD214 ForD Test Ext Rev | gcatggcgcaggtgcacacc | Confirmation of ForD deletion internal rev |
| RD215 ForL 1.1 For | gctcggttgccgccgggcgtttttaTCTAGAgcagcacgccgaccagcacg | pCRISPomyces-2 template 1 left flank ForL |
| RD216 ForL 1.2 Rev | GCTGCTGCGACCAGGCGAGCTCGCtagtcccgccatggcggacc | pCRISPomyces-2 template 1 right flank ForL |
| RD217 ForL 2.1 For | GCGAGCTCGCCTGGTCGCAGCAGCcatggacgaactccctttcgcc | pCRISPomyces-2 template 2 left flank ForL |
| RD218 ForL 2.2 Rev | gcaacgcggcctttttacggttcctggccTCTAGAgctaaggaggtggccgagg | pCRISPomyces-2 template 2 right flank ForL |
| RD219 ForL gRNA For | ACGCgagccccaactgccttggta | ForL gRNA forward |
| RD220 ForL gRNA Rev | AAACtaccaaggcagttggggctc | ForL gRNA reverse |
| RD221 ForL Test Int For | cgagggcgagcagcaggcg | Confirmation of ForL deletion external for |
| RD222 ForL Test Int Rev | cggcacgcgagttcggtggc | Confirmation of ForL deletion external rev |

| | | |
|---|---|---|
| RD223 ForL Test Ext For | cgctcggtcgccacggcc | Confirmation of FoL deletion internal for |
| RD224 ForL Test Ext Rev | ccgcgtgatgacagatgcgcc | Confirmation of ForL deletion internal rev |
| RD235 ForR 1.1 For | gctcggttgccgccgggcgtttttttaTCTAGAct tcggcaagcagttcttcg | pCRISPomyces-2 template 1 left flank ForR |
| RD236 ForR 1.2 Rev | GCTGCTGCGACCAGGCGAGCTC GCggtcatggttctccttgtcc | pCRISPomyces-2 template 1 right flank ForR |
| RD237 ForR 2.1 For | GCGAGCTCGCCTGGTCGCAGCA GCacatgacccggcaggtcgcc | pCRISPomyces-2 template 2 left flank ForR |
| RD238 ForR 2.2 Rev | gcaacgcggcctttttacggttcctggccTCTA GAccgagccgtgcgcgttgacg | pCRISPomyces-2 template 2 right flank ForR |
| RD239 ForR gRNA For | ACGCcgtagaggaactcctcggagta | ForR gRNA forward |
| RD240 ForR gRNA Rev | AAACtactccgaggagttcctctacg | ForR gRNA reverse |
| RD241 ForR Test Int For | gcacatacgccatcaggtcgc | Confirmation of ForR deletion external for |
| RD242 ForR Test Int Rev | ggtccatgcgttgggcctcg | Confirmation of ForR deletion external rev |
| RD243 ForR Test Ext For | cggcgagcgggtcttcgac | Confirmation of ForR deletion internal for |
| RD244 ForR Test Ext Rev | cgtgccggtcgttctgcttgg | Confirmation of ForR deletion internal rev |
| RD245 ForS 1.1 For | gctcggttgccgccgggcgtttttttaTCTAGAg aggagcacctcaccatgacg | pCRISPomyces-2 template 1 left flank ForS |
| RD246 ForS 1.2 Rev | GCTGCTGCGACCAGGCGAGCTC GCgctcatacggtgccctccac | pCRISPomyces-2 template 1 right flank ForS |
| RD247 ForS 2.1 For | GCGAGCTCGCCTGGTCGCAGCA GCtgatccgacccaacggggac | pCRISPomyces-2 template 2 left flank |

| | | |
|---|---|---|
| | | ForS |
| RD248 ForS 2.2 Rev | gcaacgcggcctttttacggttcctggccTCTAGAgaagcgccggtgatcatgacg | pCRISPomyces-2 template 2 right flank ForS |
| RD249 ForS gRNA For | ACGCgggtttgtggacgtaggtga | ForS gRNA forward |
| RD250 ForS gRNA Rev | AAACtcacctacgtccacaaaccc | ForS gRNA reverse |
| RD251 ForS Test Int For | cctcgtcgaggacagcacgg | Confirmation of ForS deletion external for |
| RD252 ForS Test Int Rev | cgatgcgtacgtcgacgttgc | Confirmation of ForS deletion external rev |
| RD253 ForS Test Ext For | gcctcttgcgcggcttgagc | Confirmation of ForS deletion internal for |
| RD254 ForS Test Ext Rev | cgcgtcgaagcaggagacgg | Confirmation of ForS deletion internal rev |
| RD265 ForU 1.1 For | gctcggttgccgccgggcgtttttaTCTAGAcgtagtactcgccgcggttc | pCRISPomyces-2 template 1 left flank ForU |
| RD266 ForU 1.2 Rev | GCTGCTGCGACCAGGCGAGCTCGCgatgagtgaccagcagcgcc | pCRISPomyces-2 template 1 right flank ForU |
| RD267 ForU 2.1 For | GCGAGCTCGCCTGGTCGCAGCAGCcatgtggagctgccctcactc | pCRISPomyces-2 template 2 left flank ForU |
| RD268 ForU 2.2 Rev | gcaacgcggcctttttacggttcctggccTCTAGAgcttcttgccgccgacgttg | pCRISPomyces-2 template 2 right flank ForU |
| RD269 ForU gRNA For | ACGCgtggccctcttgagctacgt | ForU gRNA forward |
| RD270 ForU gRNA Rev | AAACacgtagctcaagagggccac | ForU gRNA reverse |
| RD271 ForU Test Int For | ggtccactgcggcaggtcg | Confirmation of ForU deletion external for |
| RD272 ForU Test Int Rev | gcgcggttcctcgtcgagcg | Confirmation of ForU deletion external rev |

| | | |
|---|---|---|
| RD273 ForU Test Ext For | ggtcacgaagtcgacgtcgg | Confirmation of ForU deletion internal for |
| RD274 ForU Test Ext Rev | cggtagccgtggtaggtgagg | Confirmation of ForU deletion internal rev |
| RD307 ForD pMS82 pF | gccgagaaccTAGGATCCAAGCTTgtggagctgccctcactctc | ForD pMS82-promotor for complementation forward |
| RD308 ForD pMS82 pR | cgttctcggtgagctcgggcacggtgaggtgctcctcctg | ForD promotor-gene for complementation reverse |
| RD309 ForD pMS82 gF | caggaggagcacctcaccgtgcccgagctcaccgagaacg | ForD promotor-gene for complementation forward |
| RD310 ForD PMS82 gR | CTGGTACCATGCATAGATCTAAGCTTcctcggtggcgctcatacgg | ForD gene-pMS82 for complementation reverse |
| RD311 ForL pMS82 pF | gccgagaaccTAGGATCCAAGCTTgattcttcggcgcacgacag | ForL pMS82-promotor for complementation forward |
| RD312 ForL pMS82 pR | cgacgatcaacgtggtgtgcataccggctcccatcggttgc | ForL promotor-gene for complementation reverse |
| RD313 ForL pMS82 gF | gcaaccgatgggagccggtatgcacaccacgttgatcgtcg | ForL promotor-gene for complementation forward |
| RD314 ForL pMS82 gR | CTGGTACCATGCATAGATCTAAGCTTctactcgacggggactacgc | ForL gene-pMS82 for complementation reverse |
| RD512 ForR-ForA ErmE* F | AAAAAcatatgatgaccacgcacaccgtgc | ForRA complementation under ErmE* forward pIJ10257 |
| RD513 ForR-ForA ErmE* R | AAAAAaagctttcacctcagctccctccggtc | FoRA complementation under ErmE* reverse pIJ10257 |
| RD516 ForS ErmE* F | AAAAAcatatgatgagccaggaggagccgc | ForS complementation under ErmE* forward |

| | | |
|---|---|---|
| | | pIJ10257 |
| RD517 ForS ErmE* R | AAAAAaagctttcagaggggtgctgccgtg | ForS complementation under ErmE* reverse pIJ10257 |
| RD518 ForU-ForV ErmE* F | AAAAAcatatgatggccgagatctccgccg | ForU-V complementation under ErmE* forward pIJ10257 |
| RD519 ForU-ForV ErmE* R | AAAAAaagcttcgctctgctcctgtgccgtc | ForU-V complementation under ErmE* reverse pIJ10257 |

**Supplementary Table 4 | Growth media used in this study**

| Media | Recipe (per litre) | Water | pH |
|---|---|---|---|
| LB | 10 g tryptone<br>5 g yeast extract<br>10 g NaCl (omitted when selecting with Hygromycin)<br>+/- 20 g agar | Deionised | 7.5 |
| MS | 20 g soy flour<br>20 g mannitol<br>20 g agar | Tap | As made |
| MYM | 4 g maltose<br>4 g yeast extract<br>10 g malt extract<br>18g agar | 50:50<br>Tap:Deisonised | 7.3 |

**Supplementary Table 5 | Antibiotics (and concentrations) used in this study.**

| Antibiotic | Final concentration used for selection (μg/ml) |
|---|---|
| Apramycin | 50 |
| Hygromycin | 50 |
| Kanamycin | 50 |
| Chloramphenicol | 30 |
| Nalidixic Acid | 25 |

**Supplementary Note 1 | Compound structure elucidation data for 6-chlorogenistein**

An isomer of 6-chlorogenistein – identified based on based on UV and MS characteristics, and likely to be a chlorination regioisomer - was identified in the LCMS trace but was not isolated due to low levels of production. The NMR data for 6-chlorogenistein has been reported previously[6], and is in accordance with the data reported here.

**Supplementary Figure 11 | Chemical structure of 6-chlorogenistein**

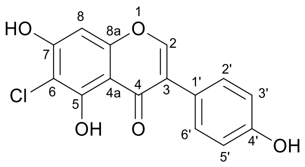


Molecular formula: $C_{15}H_9O_5Cl$

Isolated yield: 2 mg

UV (PDA): $\lambda_{max}$ = 259 nm

HRMS (ESI) *m/z*: calculated $[M + H]^+$ = 305.0211; observed $[M + H]^+$ = 305.0212, Δ = 0.33 ppm

**Supplementary Table 6 | NMR data for 6-chlorogenistein in $CD_3OD$ at 400 MHz for $^1H$ and 100 MHz for $^{13}C$.**

| Position | $\delta_C$ ppm | $\delta_H$ ppm (no. of protons, multiplicity, J in Hz) | HMBC |
|---|---|---|---|
| 2 | 155.0 | 8.2 | |
| 3 | 123.1 | | 2, 2’, 6’ |
| 4 | 182.3 | | 2 |
| 4a | 107.0 | | 8 |
| 5 | 162.2 | | |
| 6 | 99.5 | | 8 |
| 7 | 162.0 | | 8 |
| 8 | 100.5 | 6.4 | |
| 8a | 155.0 | | 2 |
| 1’ | 125.2 | | 2, 3’, 5’ |
| 2’ | 116.5 | 6.8 | 6’ |
| 3’ | 131.6 | 7.4 | 5’ |
| 4’ | 159.2 | | 2’, 3’, 5’, 6’ |

| | | | |
|---|---|---|---|
| 5’ | 131.6 | 7.4 | 3’ |
| 6’ | 116.5 | 6.9 | 2’ |

**Supplementary Figure 12 | $^{1}H$ NMR spectrum ($CD_3OD$, 400 MHz) for 6-chlorogenistein.**

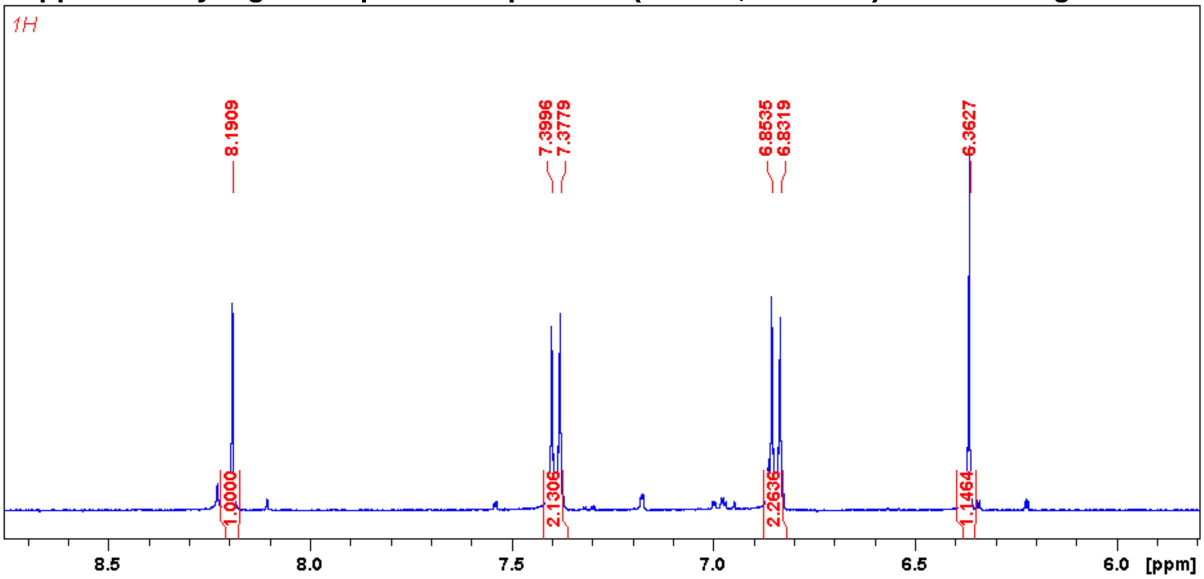


**Supplementary Figure 13 | $^{13}C$ NMR spectrum ($CD_3OD$, 100 MHz) for 6-chlorogenistein.**

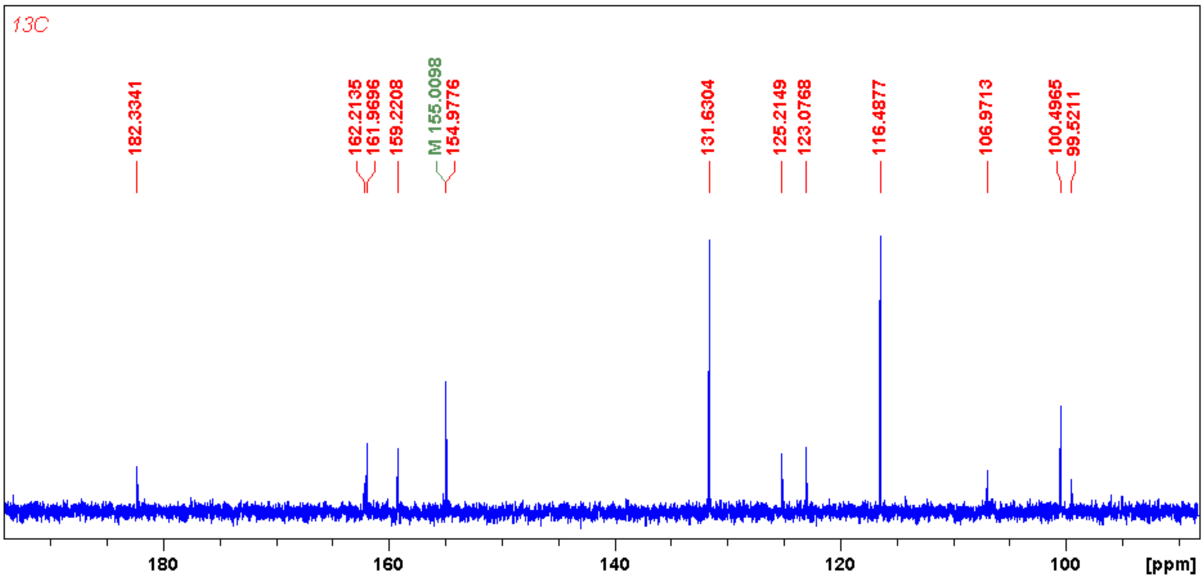

**Supplementary Figure 14 | HSQC spectrum ($CD_3OD$) for 6-chlorogenistein.**

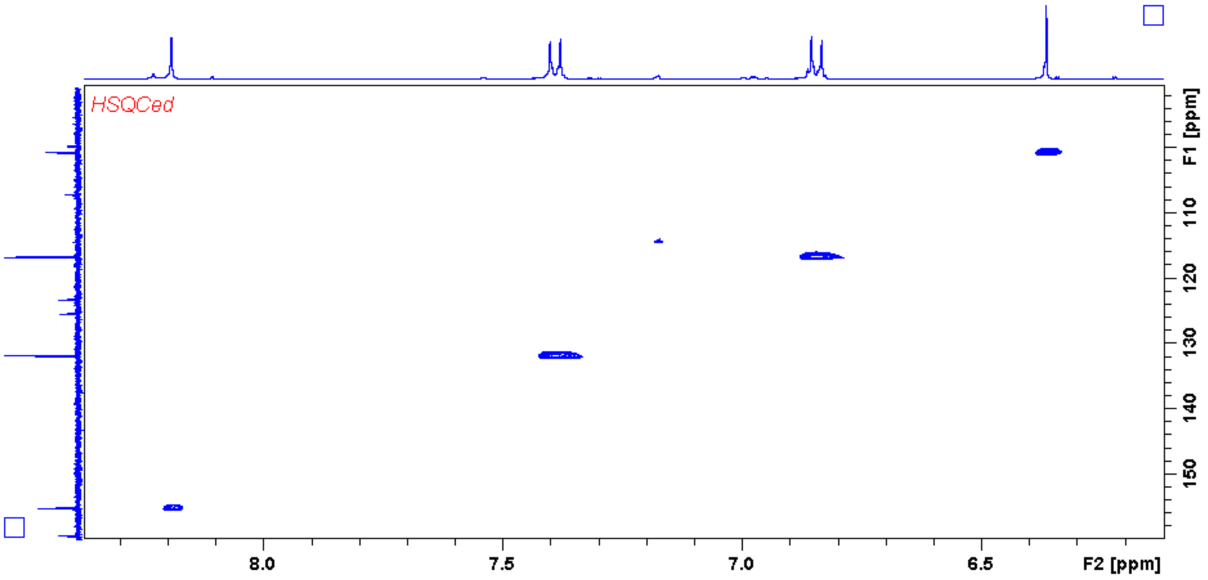


**Supplementary Figure 15 | HMBC spectrum ($CD_3OD$) for 6-chlorogenistein.**

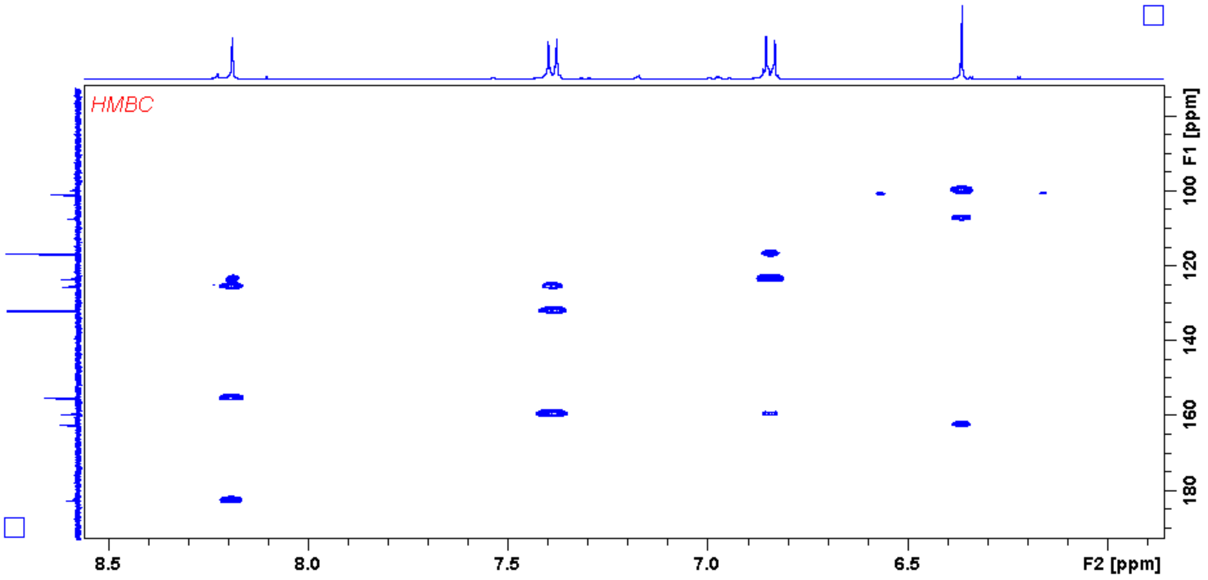

**Supplementary Note 2 | Formicapyridine A-F (1-6) and fasamycin F (13) structure elucidation data.**

**COMPOUND 1 (formicapyridine A)**

**Supplementary Figure 16 | Chemical structure of compound 1**

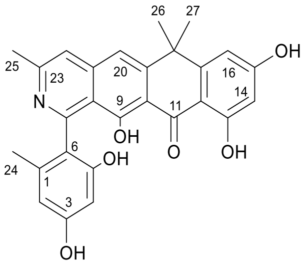


Molecular formula: $C_{27}H_{23}NO_6$

Isolated yield: 1 mg

UV (PDA): $\lambda_{max}$ = 228, 249, 272, and 391 nm

Optical activity: $[\alpha]_D^{20}$ = +9.890

HRMS (ESI) *m/z*: calculated $[M + H]^+$ = 458.1598; observed $[M + H]^+$ = 458.1600, Δ = 0.44 ppm

**Supplementary Table 7 | NMR data for compound 1 in $CD_3OD$ at 400 MHz for $^1H$ and 100 MHz for $^{13}C$.**

| Position | $\delta_C$ ppm | $\delta_H$ ppm (no. of protons, multiplicity, J in Hz) | HMBC | NOESY |
|---|---|---|---|---|
| 1 | 138.8 | | 24 | |
| 2 | 109.4 | 6.3 (1H, d, 1.64) | 24 | 24 |
| 3 | not detected | | | |
| 4 | 101.2 | 6.3 (1H, d, 1.64) | | |
| 5 | 157.0 | | | |
| 6 | 120.9 | | 2, 4, 24 | |
| 7 | 159.9 | | | |
| 8 | 118.6 | | 20, 22 | |
| 9 | 167.5 | | | |
| 10 | 111.3 | | 20 | |
| 11 | 191.6 | | | |
| 12 | 108.5 | | 14 | |
| 13 | 167.7 | | 14 | |

| | | | | |
|---|---|---|---|---|
| 14 | 102.5 | 6.3 (1H, d, 2.21) | 16 | |
| 15 | 167.8 | | | |
| 16 | 107.9 | 6.7 (1H, d, 2.21) | 14 | 26, 27 |
| 17 | 155.7 | | 26, 27 | |
| 18 | 40.7 | | 20, 26, 27 | |
| 19 | 155.1 | | 26, 27 | |
| 20 | 115.7 | 7.7 (1H, s) | 22 | 26, 27 |
| 21 | 144.7 | | | |
| 22 | 121.3 | 7.7 (1H, s) | 20, 25 | 25 |
| 23 | 152.6 | | 25 | |
| 24 | 20.2 | 1.9 (3H, s) | | 2 |
| 25 | 22.1 | 2.7 (3H, s) | | 6 |
| 26 | 34.5 | 1.8 (3H, s) | 27 | 16, 20 |
| 27 | 34.7 | 1.8 (3H, s) | 26 | 16, 20 |

**Supplementary Figure 17 | $^{1}H$ NMR spectrum ($CD_3OD$, 400 MHz) for compound 1.**

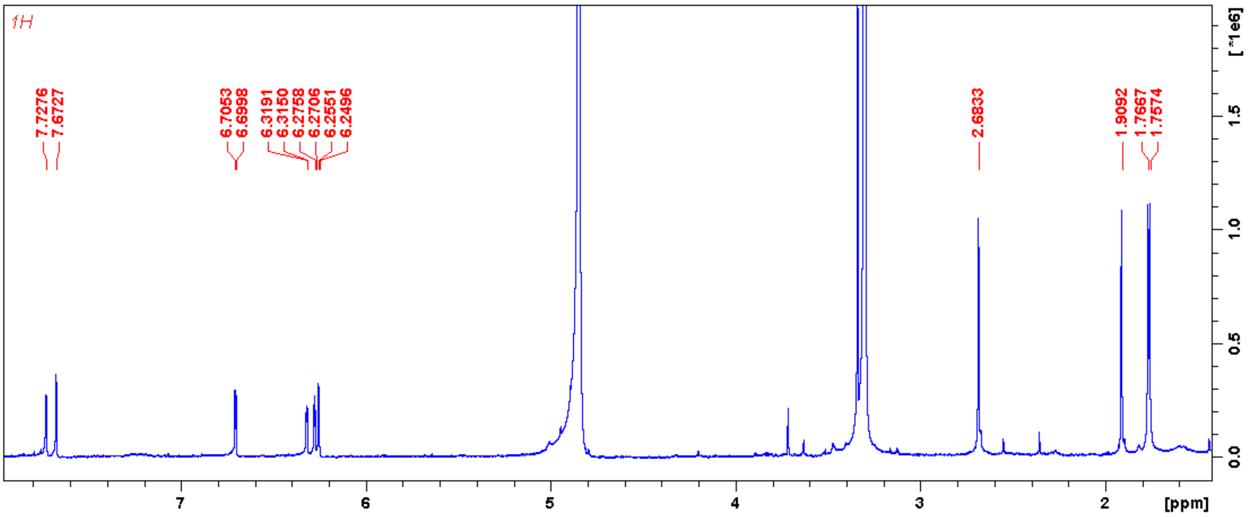

**Supplementary Figure 18 | $^{13}$C NMR spectrum ($CD_3OD$, 100 MHz) for compound 1.**

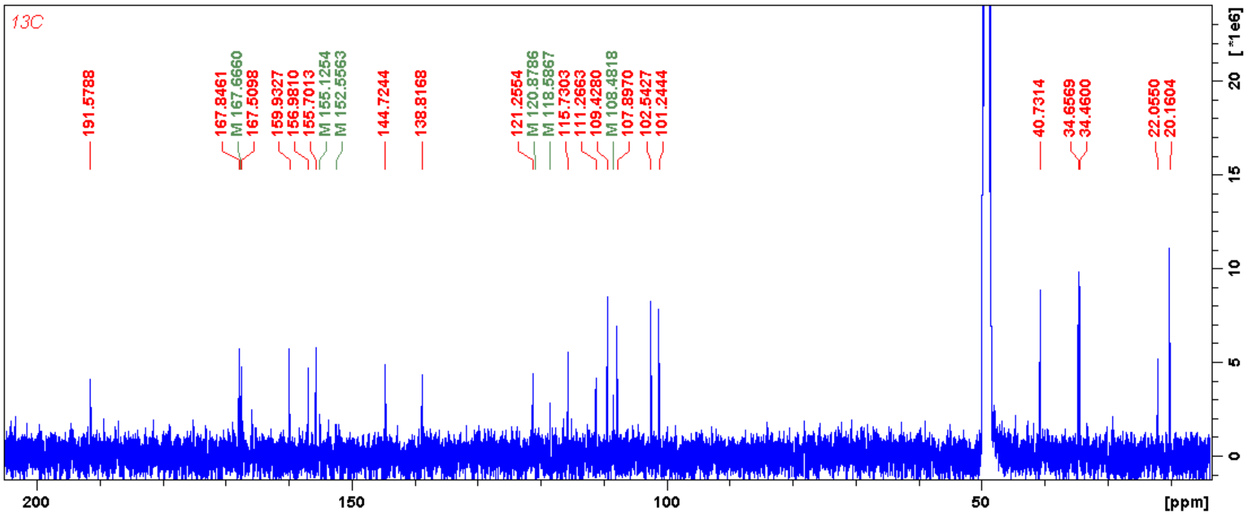


**Supplementary Figure 19 | HSQC spectrum ($CD_3OD$) for compound 1.**

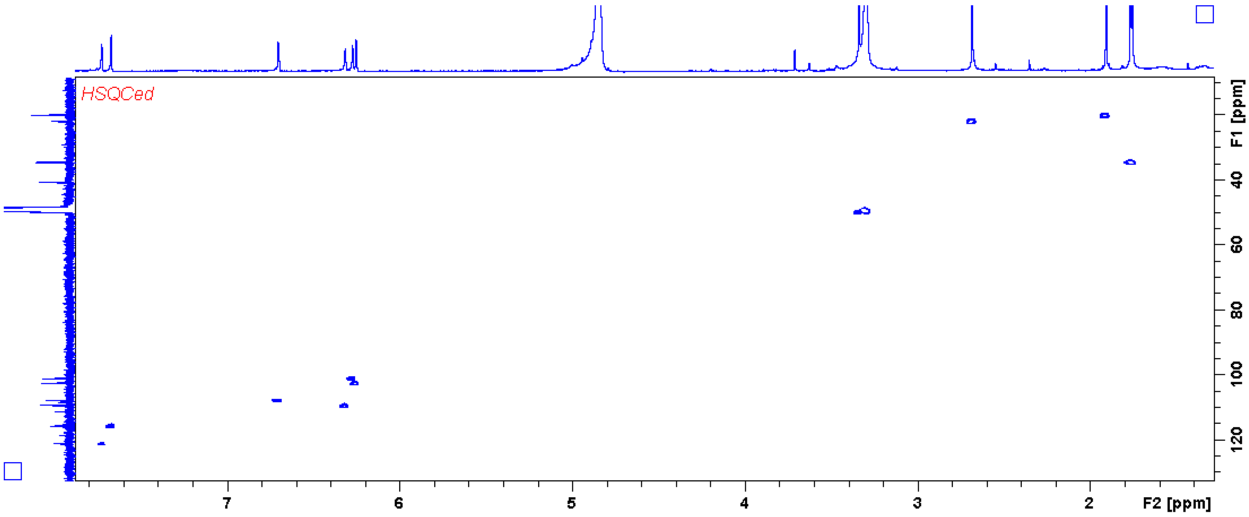


**Supplementary Figure 20 | HMBC spectrum ($CD_3OD$) for compound 1.**

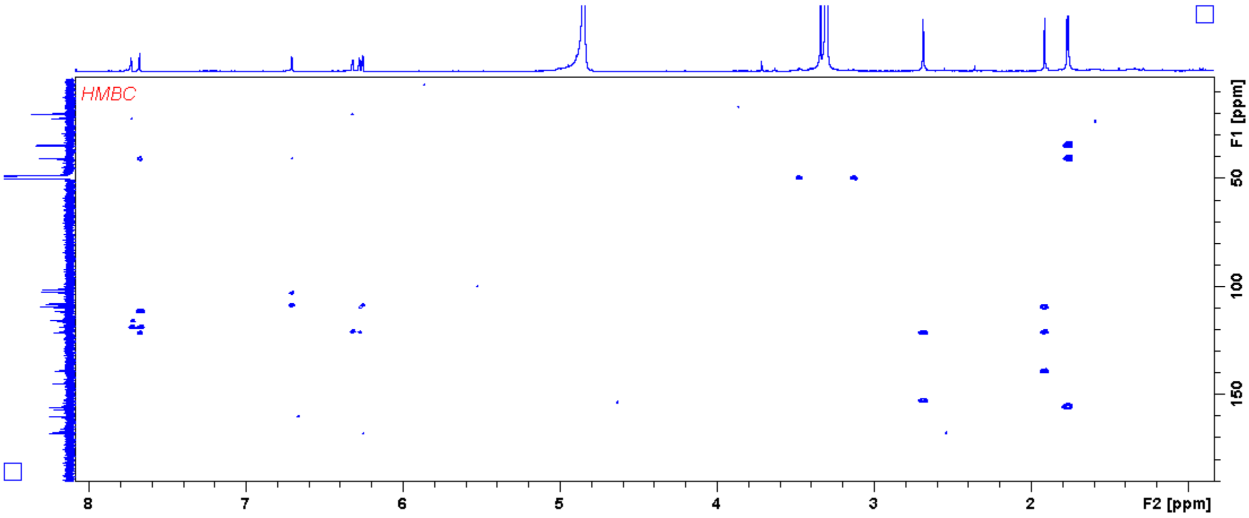

**Supplementary Figure 20 | NOESY spectrum ($CD_3OD$) for compound 1.**

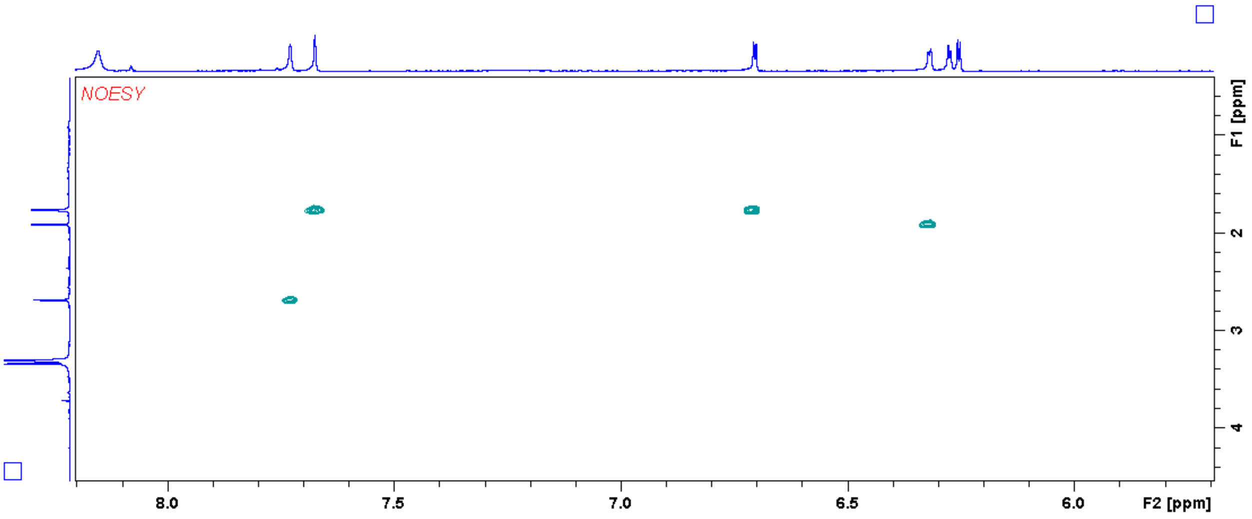


**COMPOUND 2 (formicapyridine B)**

**Supplementary Figure 22 | Chemical structure of compound 2**

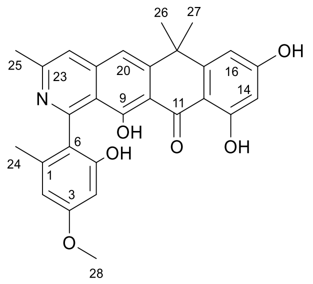


Molecular formula: $C_{28}H_{25}NO_6$

Isolated yield: 1 mg

UV (PDA): $\lambda_{max}$ = 229, 249, 272, and 392 nm

Optical activity: $[\alpha]_D^{20}$ = +7.692

HRMS (ESI) *m/z*: calculated $[M + H]^+$ = 472.1755; observed $[M + H]^+$ = 472.1753, Δ = -0.42 ppm.

**Supplementary Table 8 | NMR data for compound 2 in $CD_3OD$ at 400 MHz for $^1H$ and 100 MHz for $^{13}C$.**

| Position | $\delta_C$ ppm | $\delta_H$ ppm (no. of protons, multiplicity, J in Hz) | $^1H$ – $^1H$ COSY | HMBC | NOESY |
|---|---|---|---|---|---|
| 1 | 138.6 | | | 24 | |
| 2 | 107.7 | 6.4 (1H, d, 2.27) | 4 | 4, 24 | 24, 28 |

| | | | | | |
|---|---|---|---|---|---|
| 3 | 161.7 | | | 4, 28 | |
| 4 | 99.9 | 6.3 (1H, d, 2.27) | 2 | 2 | 28 |
| 5 | 156.7 | | | 4 | |
| 6 | 124.9 | | | 2, 4, 24 | |
| 7 | 160.1 | | | | |
| 8 | 118.6 | | | 20, 22 | |
| 9 | 167.3 | | | | |
| 10 | 110.6 | | | 20 | |
| 11 | 191.8 | | | | |
| 12 | 108.6 | | | 14, 16 | |
| 13 | 167.5 | | | 14 | |
| 14 | 102.5 | 6.2 (1H, d, 2.25) | 16 | 16 | |
| 15 | 167.4 | | | 14, 16 | |
| 16 | 107.6 | 6.7 (1H, d, 2.25) | 14 | 14 | 26, 27 |
| 17 | 155.8 | | | 26, 27 | |
| 18 | 40.5 | | | 16, 20, 26, 27 | |
| 19 | 152.7 | | | 26, 27 | |
| 20 | 115.6 | 7.6 (1H, s) | | 22 | 26, 27 |
| 21 | 144.4 | | | 20 | |
| 22 | 119.9 | 7.6 (1H, s) | | 20, 25 | 25 |
| 23 | 155.5 | | | 22, 25 | |
| 24 | 20.4 | 1.9 (3H, s) | | 2 | 2 |
| 25 | 23.6 | 2.7 (3H, s) | | 22 | 22 |
| 26 | 34.6 | 1.8 (3H, s) | | 27 | 16. 20 |
| 27 | 34.8 | 1.7 (3H, s) | | 26 | 16, 20 |
| 28 | 55.8 | 3.8 (3H, s) | | | 2, 4 |

**Supplementary Figure 23 | $^{1}H$ NMR spectrum ($CD_3OD$, 400 MHz) for compound 2.**

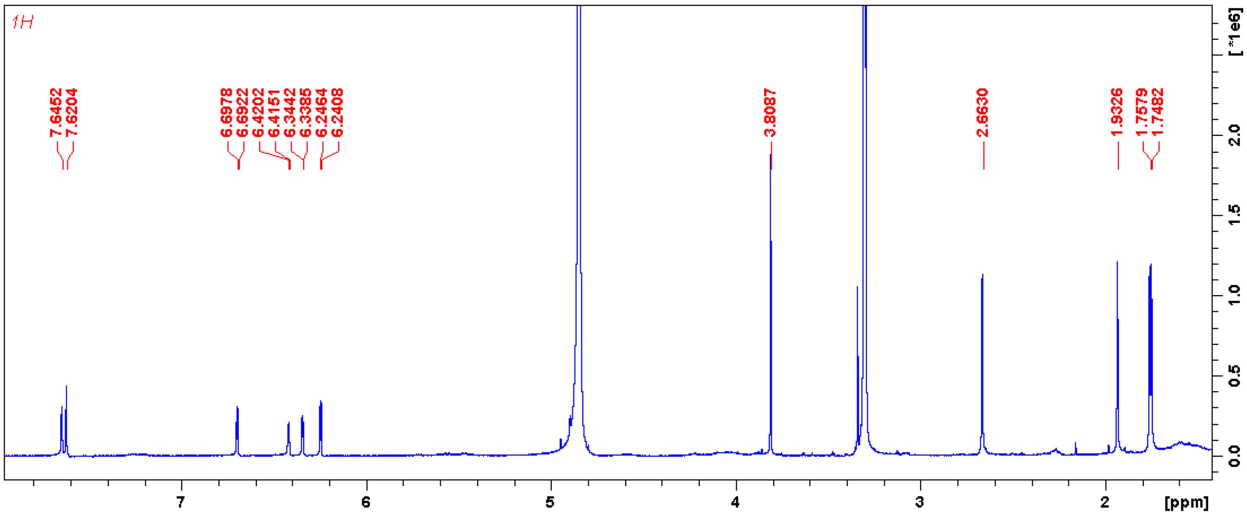


**Supplementary Figure 24 | $^{13}C$ NMR spectrum ($CD_3OD$, 100 MHz) for compound 2.**

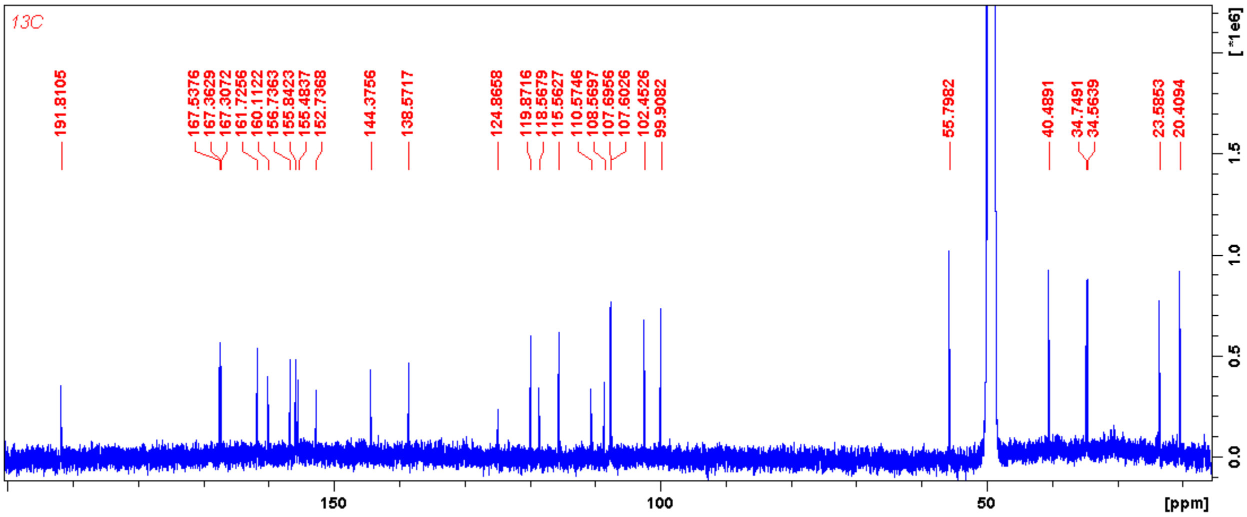


**Supplementary Figure 25 | HSQC spectrum ($CD_3OD$) for compound 2.**

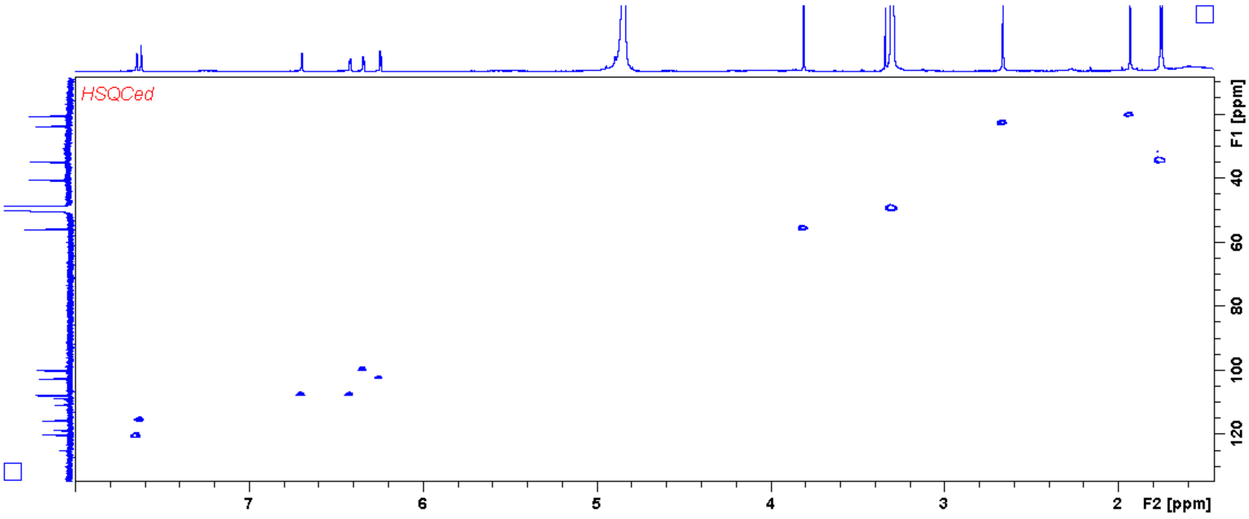

**Supplementary Figure 26 | HMBC spectrum ($CD_3OD$) for compound 2.**

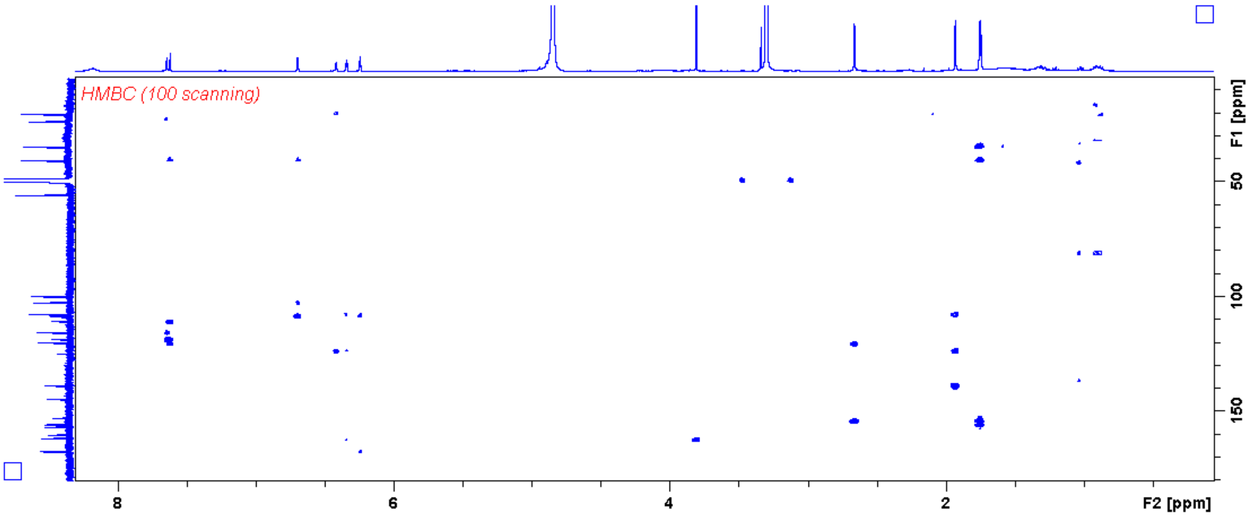


**Supplementary Figure 27 | NOESY spectrum ($CD_3OD$) for compound 2.**

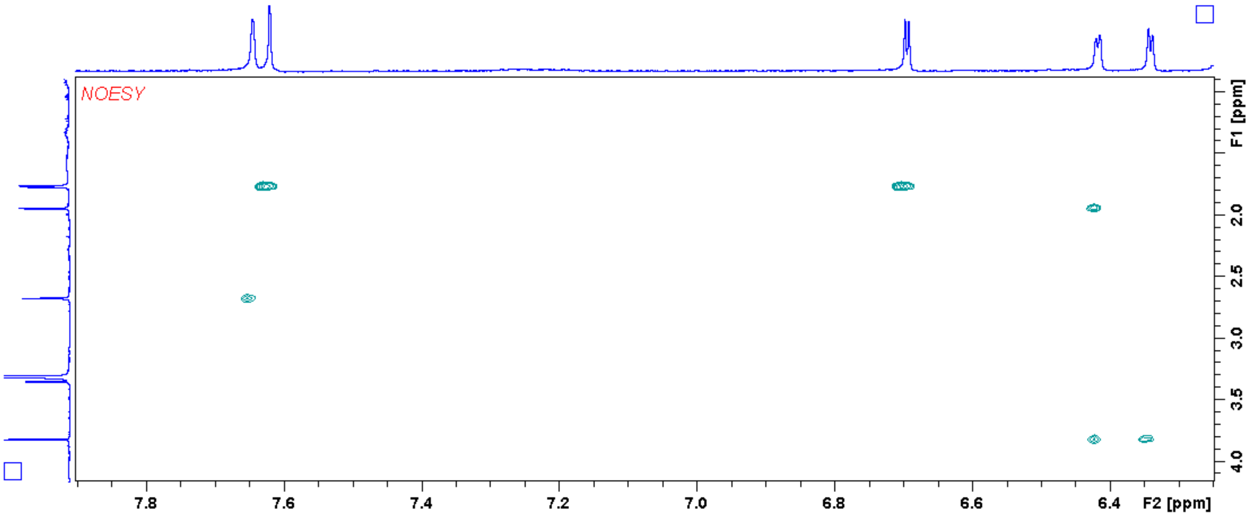


**COMPOUND 3 (formicapyridine C)**

**Supplementary Figure 28 | Chemical structure of compound 3**

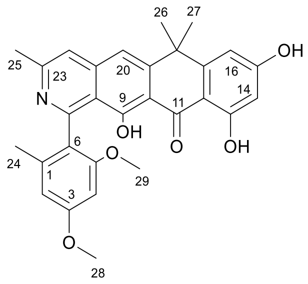

Molecular formula: $C_{29}H_{27}NO_6$

Isolated yield: 2 mg

UV (PDA): $\lambda_{max}$ = 228, 249, 272, and 391 nm

Optical activity: $[\alpha]_D^{20}$ = +8.791

HRMS (ESI) *m/z*: calculated $[M + H]^+$ = 486.1911; observed $[M + H]^+$ = 486.1905, Δ = -1.23 ppm.

**Supplementary Table 9 | NMR data for compound 3 in $CD_3OD$ at 400 MHz for $^1H$ and 100 MHz for $^{13}C$.**

| Position | $\delta_C$ ppm | $\delta_H$ ppm (no. of protons, multiplicity, J in Hz) | HMBC | NOESY |
|---|---|---|---|---|
| 1 | 138.5 | | 25 | |
| 2 | 107.7 | 6.5 (1H, d, 2.33) | 4, 24 | 28 |
| 3 | 162.4 | | 4, 28 | |
| 4 | 97.0 | 6.5 (1H, d, 2.33) | 2 | 28, 29 |
| 5 | 159.7 | | 4, 29 | |
| 6 | 125.1 | | 4, 25 | |
| 7 | 161.0 | | | |
| 8 | 118.4 | | 20 | |
| 9 | 167.2 | | | |
| 10 | 110.8 | | 20 | |
| 11 | 191.9 | | | |
| 12 | 108.5 | | 14, 16 | |
| 13 | 167.4 | | 14 | |
| 14 | 102.5 | 6.2 (1H, d, 2.30) | 16 | |
| 15 | 167.7 | | 16 | |
| 16 | 107.8 | 6.7 (1H, d, 2.30) | | 26, 27 |
| 17 | 155.8 | | 26, 27 | |
| 18 | 40.6 | | 16, 20, 26, 27 | |
| 19 | 153.6 | | 26, 27 | |
| 20 | 115.7 | 7.6 (1H, s) | 22 | 26, 27 |
| 21 | 144.4 | | 20 | |
| 22 | 120.4 | 7.6 (1H, s) | 20, 25 | 25 |
| 23 | 154.5 | | 22, 25 | |
| 24 | 20.2 | 1.9 (3H, s) | 2 | 2 |
| 25 | 23.1 | 2.6 (3H, s) | 22 | 22 |
| 26 | 34.6 | 1.7 (3H, s) | 27 | 16, 20 |

| | | | | |
|---|---|---|---|---|
| 27 | 34.6 | 1.8 (3H, s) | 26 | 16, 20 |
| 28 | 56.0 | 3.9 (3H, s) | | 2, 4 |
| 29 | 56.3 | 3.6 (3H, s) | | 4 |

**Supplementary Figure 29 | [1]H NMR spectrum ($CD_3OD$, 400 MHz) for compound 3.**

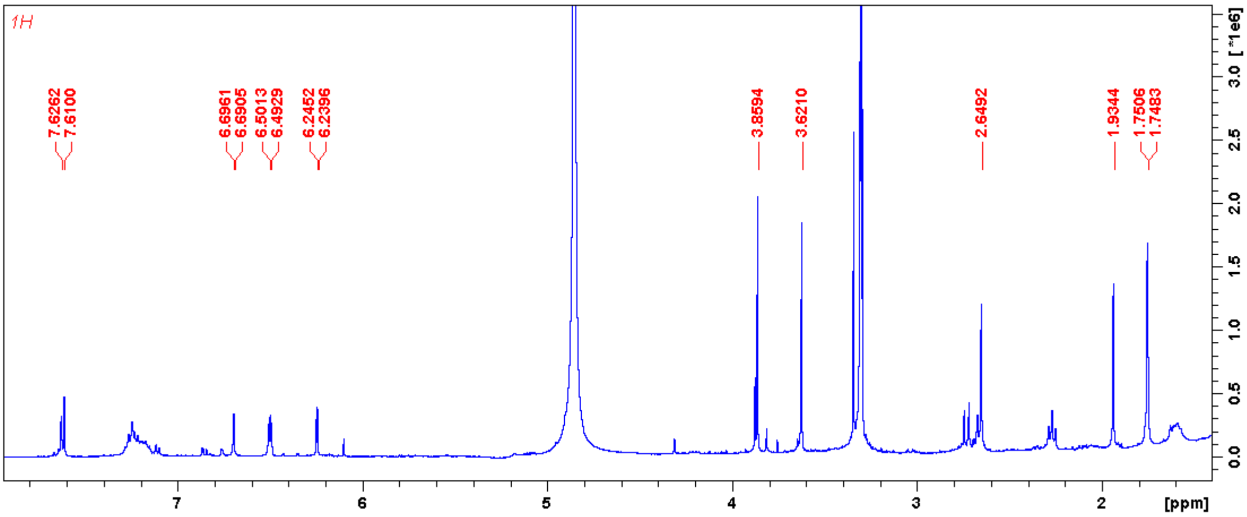

**Supplementary Figure 30 | $^{13}$C NMR spectrum ($CD_3OD$, 100 MHz) for compound 3.**

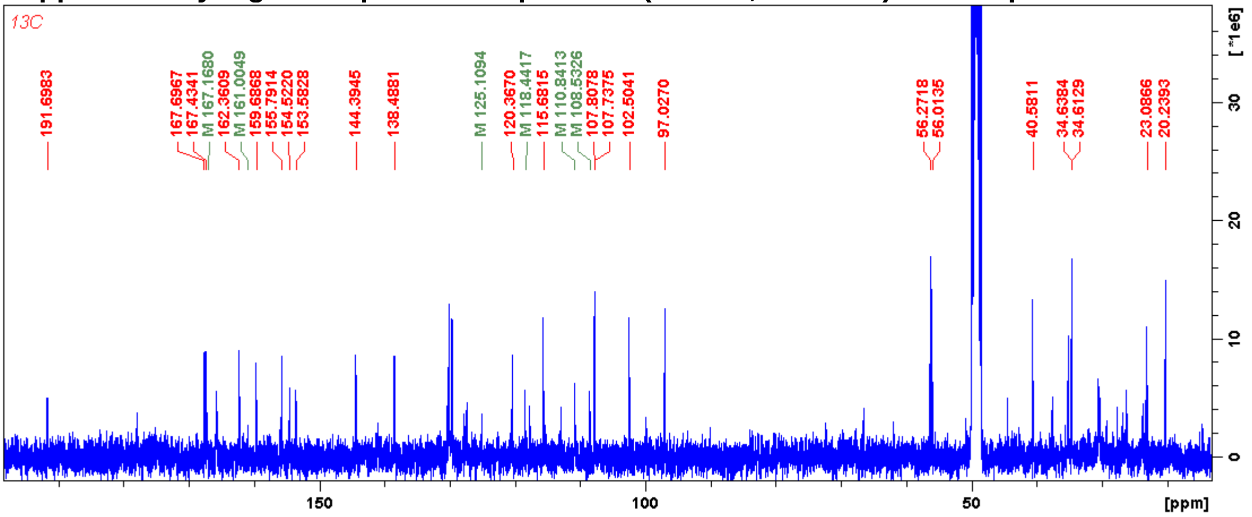


**Supplementary Figure 31 | HSQC spectrum ($CD_3OD$) for compound 3.**

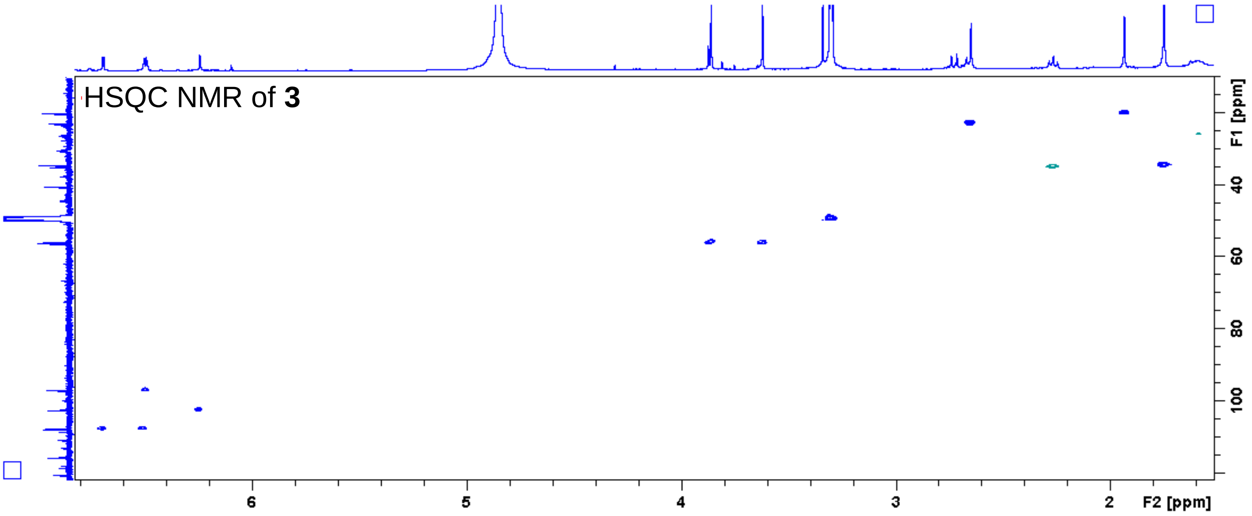


**Supplementary Figure 32 | HMBC spectrum ($CD_3OD$) for compound 3.**

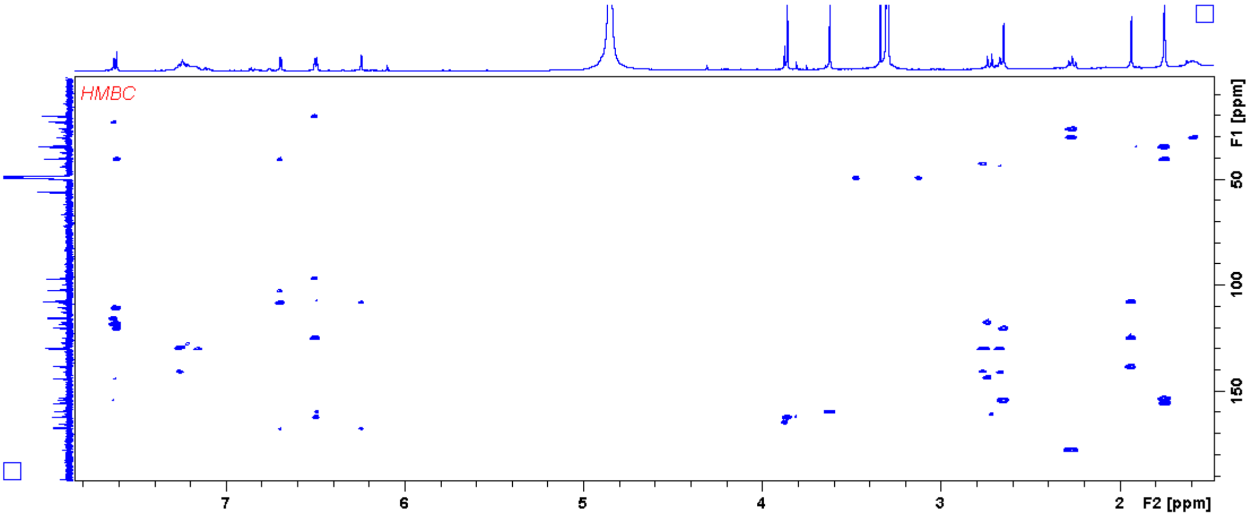

**Supplementary Figure 33 | NOESY spectrum ($CD_3OD$) for compound 3.**

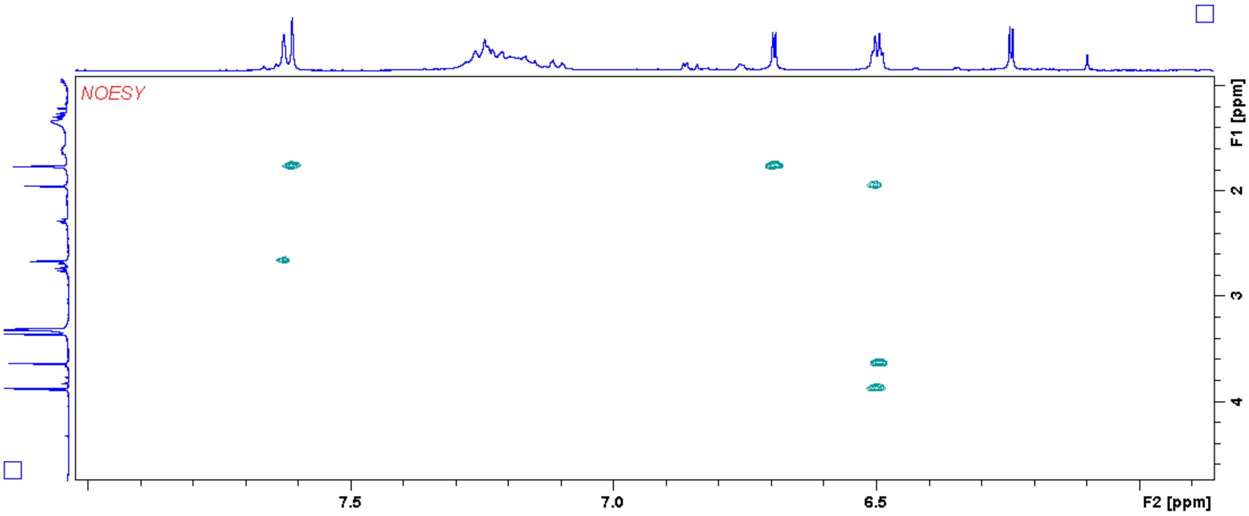


**COMPOUND 4 (formicapyridine D)**

**Supplementary Figure 34 | Chemical structure of compound 4**

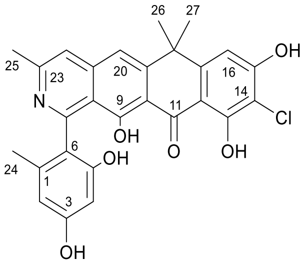


Molecular formula: $C_{27}H_{22}NO_6Cl$

Isolated yield: 0.7 mg

UV (PDA): $\lambda_{max}$ = 228, 252, and 391 nm

Optical activity: $[\alpha]_D^{20}$ = +10.989

HRMS (ESI) *m/z*: calculated $[M + H]^+$ = 492.1218; observed $[M + H]^+$ = 492.1208, Δ = 2.03 ppm.

**Supplementary Table 10 | NMR data for compound 4 in $CD_3OD$ at 400 MHz for $^1H$.**

| Position | $\delta_H$ ppm (no. of protons, multiplicity, J in Hz) | NOESY |
|---|---|---|
| 2 | 6.2929 (1H, d, 2.32) | 24 |
| 4 | 6.2511 (1H, d, 2.32) | |
| 16 | 6.8255 (1H, s) | 26, 27 |

| | | |
|---|---|---|
| 20 | 7.5921 (1H, s) | 26, 27 |
| 22 | 7.5921 (1H, s) | 25 |
| 24 | 1.8789 (3H, s) | 2 |
| 25 | 2.6475 (3H, s) | 22 |
| 26 | 1.7456 (3H, s) | 16, 20 |
| 27 | 1.7456 (3H, s) | 16, 20 |

**Supplementary Figure 35 | $^1H$ NMR spectrum ($CD_3OD$, 400 MHz) for compound 4.**

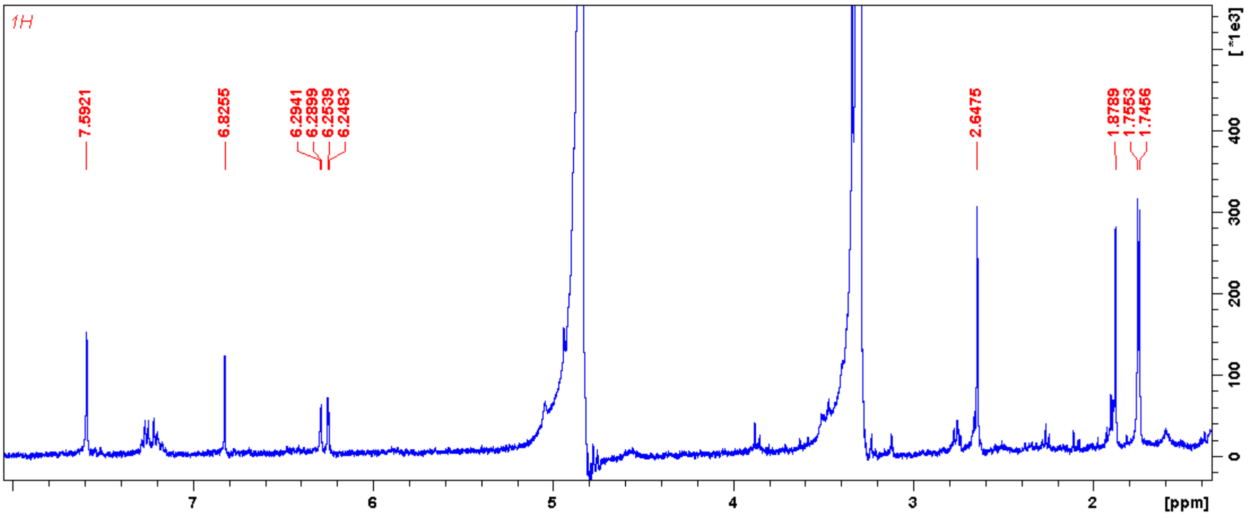


**Supplementary Figure 36 | NOESY spectrum ($CD_3OD$) for compound 4.**

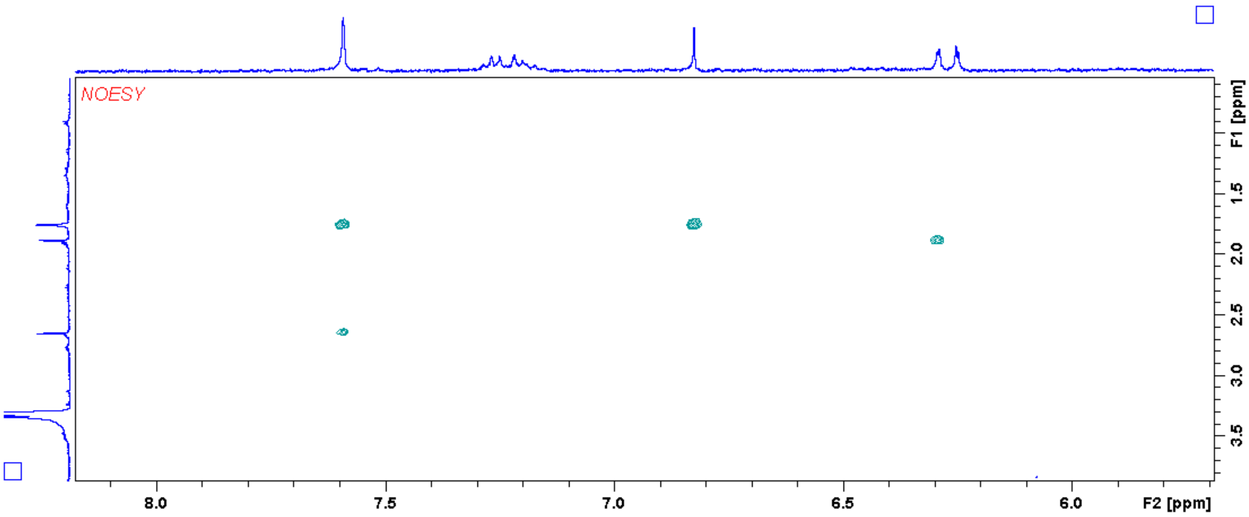

**COMPOUND 5 (formicapyridine E)**

**Supplementary Figure 37 | Chemical structure of compound 5**

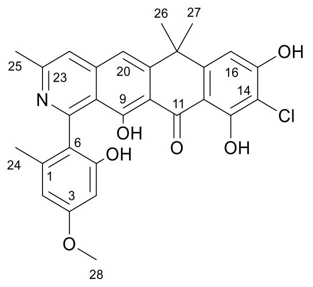


Molecular formula: $C_{28}H_{24}NO_6Cl$

Isolated yield: 1 mg

UV (PDA): $\lambda_{max}$ = 218, 252, and 391 nm

Optical activity: $[\alpha]_D^{20}$ = +12.088

HRMS (ESI) *m/z*: calculated $[M + H]^+$ = 506.1365; observed $[M + H]^+$ = 506.1362, Δ = -0.59 ppm.

**Supplementary Table 11 | NMR data for compound 5 in $CD_3OD$ at 400 MHz for $^1H$ and 100 MHz for $^{13}C$.**

| Position | $\delta_C$ ppm | $\delta_H$ ppm (no. of protons, multiplicity, J in Hz) | HMBC | NOESY |
|---|---|---|---|---|
| 1 | 138.8 | | 24 | |
| 2 | 107.8 | 6.4 (1H, d, 2.22) | 24 | 24, 28 |
| 3 | 162.4 | | 28 | |
| 4 | 99.9 | 6.3 (1H, d, 2.22) | | 28 |
| 5 | 156.9 | | | |
| 6 | 122.7 | | 2, 24 | |
| 7 | 159.9 | | | |
| 8 | 118.6 | | 20, 22 | |
| 9 | 162.3 | | | |
| 10 | 111.0 | | 20 | |
| 11 | 191.6 | | | |
| 12 | 109.0 | | 16 | |
| 13 | 165.2 | | | |
| 14 | 97.3 | | | |
| 15 | 163.0 | | | |

| | | | | |
|---|---|---|---|---|
| 16 | 107.5 | 6.8 (1H, s) | | 26, 27 |
| 17 | 154.1 | | 26, 27 | |
| 18 | 40.5 | | 16, 20, 26, 27 | |
| 19 | 153.1 | | 26, 27 | |
| 20 | 115.9 | 7.7 (1H, s) | 22 | 26, 27 |
| 21 | 144.8 | | | |
| 22 | 121.0 | 7.7 (1H, s) | 20, 25 | 25 |
| 23 | 153.7 | | 25 | |
| 24 | 20.3 | 1.9 (3H, s) | 22 | 2 |
| 25 | 22.5 | 2.7 (3H, s) | | 22 |
| 26 | 34.4 | 1.7 (3H, s) | 27 | 16, 20 |
| 27 | 34.6 | 1.7 (3H, s) | 26 | 16, 20 |
| 28 | 55.9 | 3.8 (3H, s) | | 2, 4 |

**Supplementary Figure 38 | $^1$H NMR spectrum ($CD_3OD$, 400 MHz) for compound 5.**

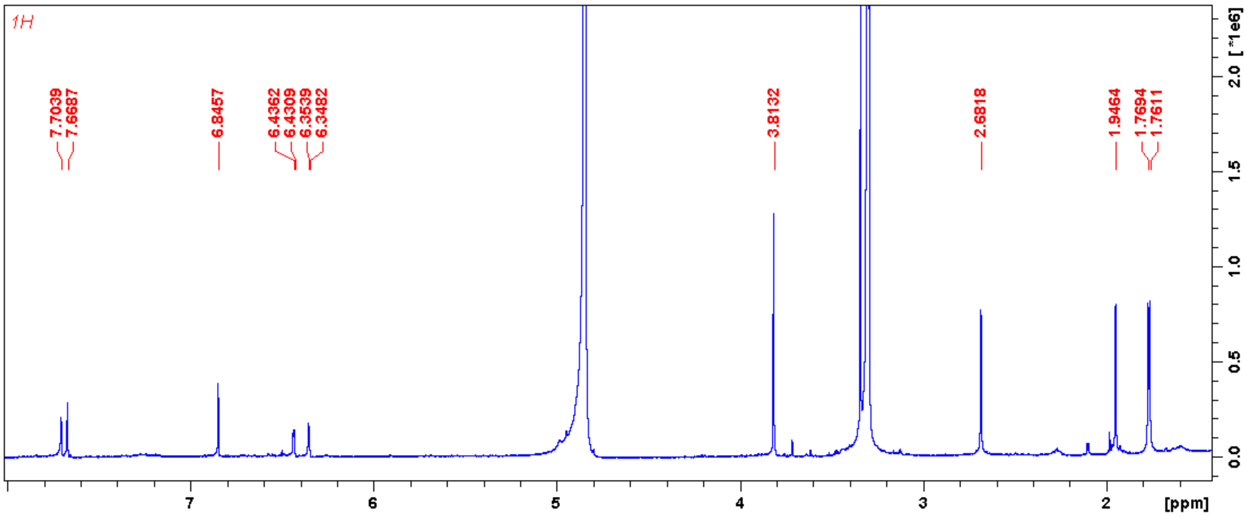

**Supplementary Figure 39 | $^{13}$C NMR spectrum ($CD_3OD$, 100 MHz) for compound 5.**

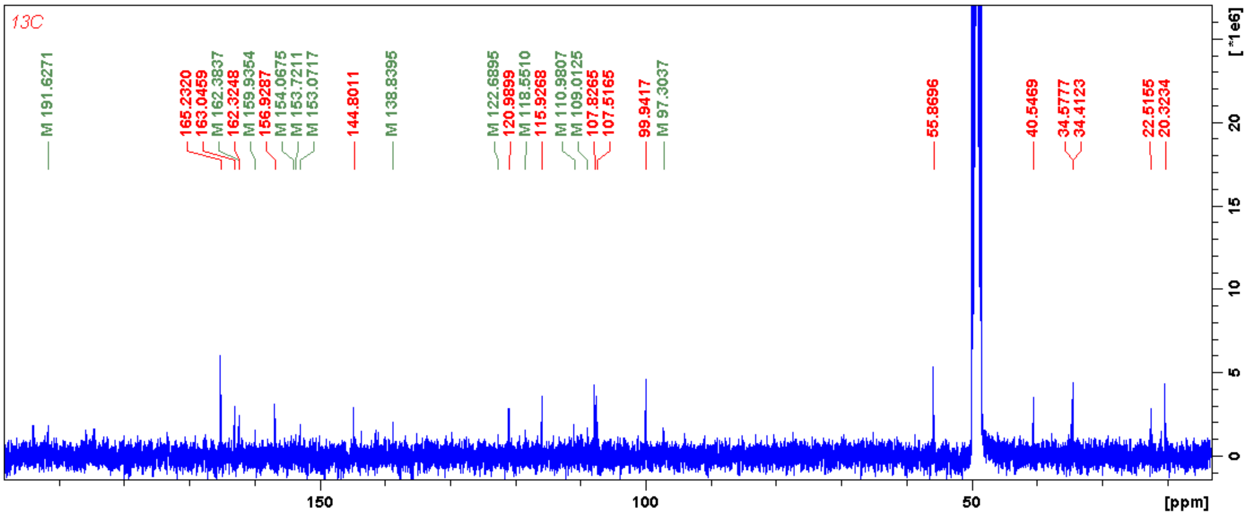


**Supplementary Figure 40 | HSQC spectrum ($CD_3OD$) for compound 5.**

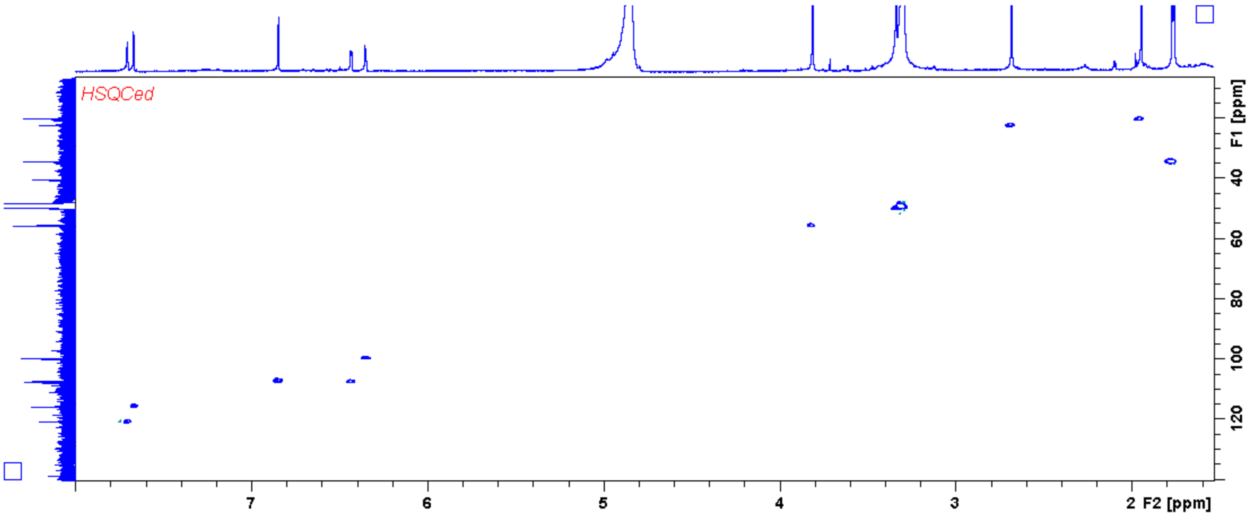


**Supplementary Figure 41 | HMBC spectrum ($CD_3OD$) for compound 5.**

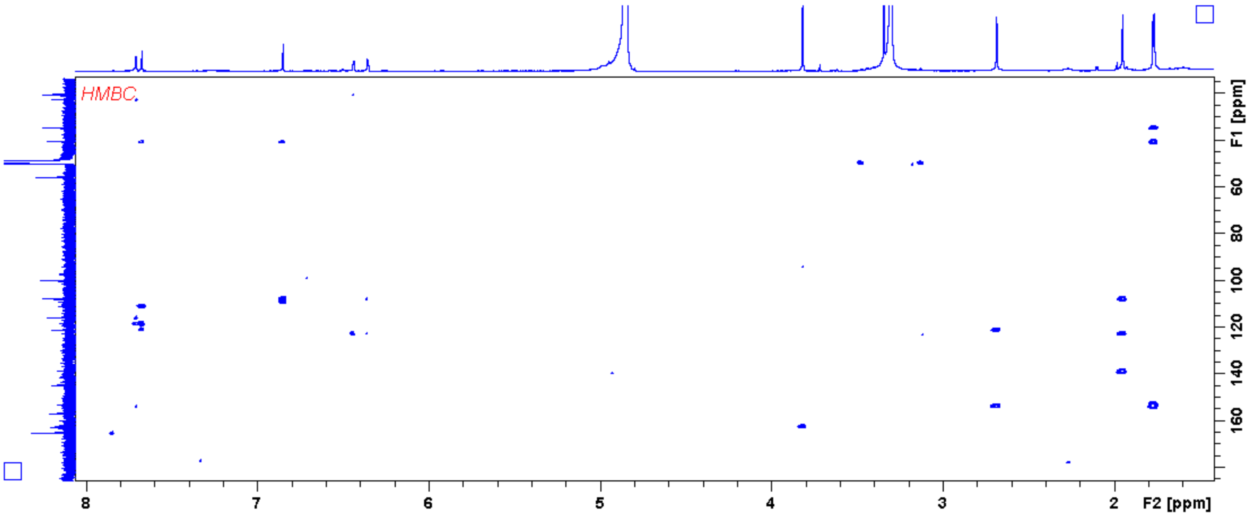

**Supplementary Figure 42 | NOESY spectrum ($CD_3OD$) for compound 5.**

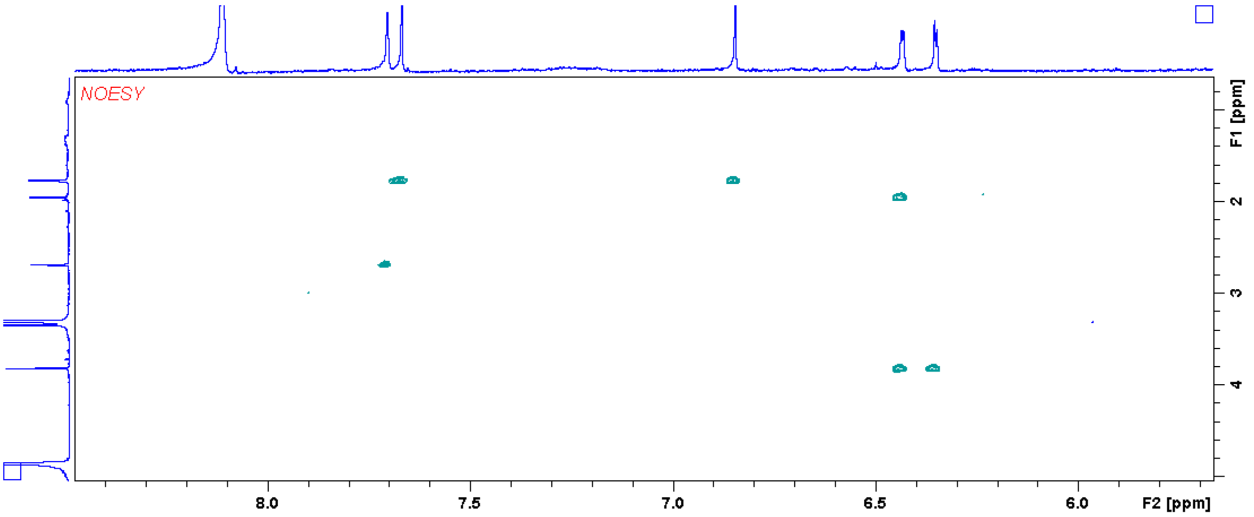


**COMPOUND 6 (formicapyridine F)**

**Supplementary Figure 43 | Chemical structure of compound 6**

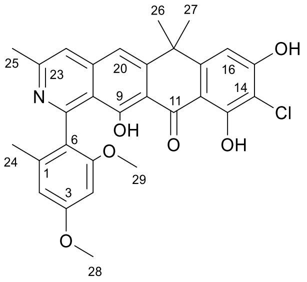


Molecular formula: $C_{29}H_{26}NO_6Cl$

Isolated yield: 0.6 mg

UV (PDA): $\lambda_{max}$ = 231, 252, and 391 nm

Optical activity: $[\alpha]_D^{20}$ = +12.821

HRMS (ESI) *m/z*: calculated $[M + H]^+$ = 520.1521; observed $[M + H]^+$ = 520.1525, Δ = 0.77 ppm.

**Supplementary Table 12 | NMR data for compound 6 in $CD_3OD$ at 400 MHz for $^1H$.**

| Position | $\delta_H$ ppm (no. of protons, multiplicity, J in Hz) | NOESY |
|---|---|---|
| 2 | 6.4915 (1H, d, 2.33) | 24 |
| 4 | 6.4834 (1H, d, 2.33) | 29 |
| 16 | 6.833 | 26, 27 |

| | | |
|---|---|---|
| 20 | 7.5951 | 26, 27 |
| 22 | 7.5863 | 25 |
| 24 | 1.9243 (3H, s) | 2 |
| 25 | 2.6384 (3H, s) | 22 |
| 26 | 1.7491 (3H, s) | 16, 20 |
| 27 | 1.7491 (3H, s) | 16, 20 |
| 28 | 3.8568 (3H, s) | 2 |
| 29 | 3.6116 (3H, s) | 4 |

**Supplementary Figure 44 | $^1$H NMR spectrum ($CD_3OD$, 400 MHz) for compound 6.**

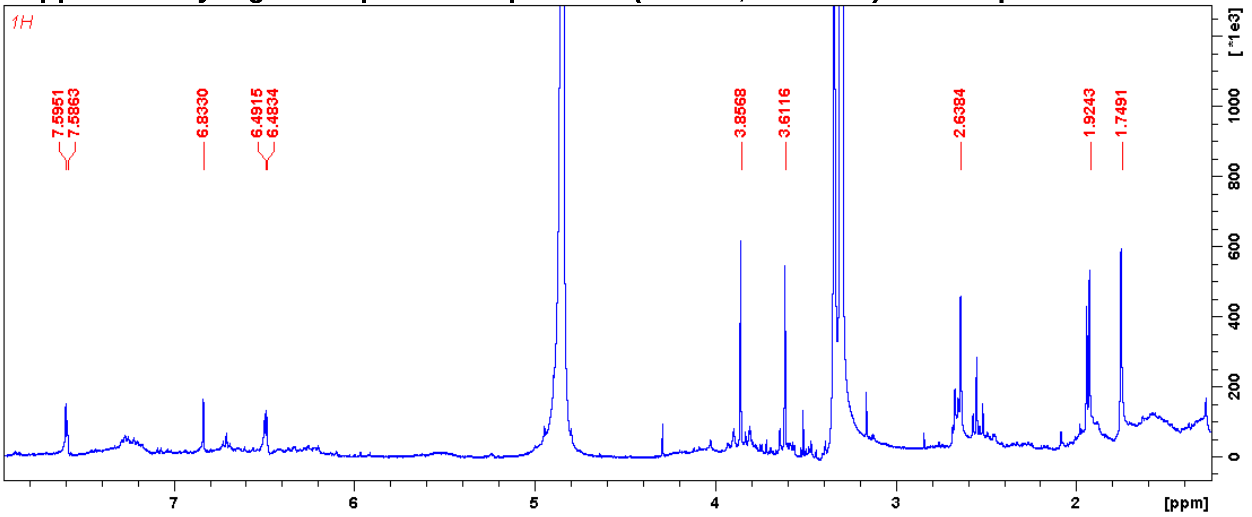


**Supplementary Figure 45 | NOESY spectrum ($CD_3OD$) for compound 6.**

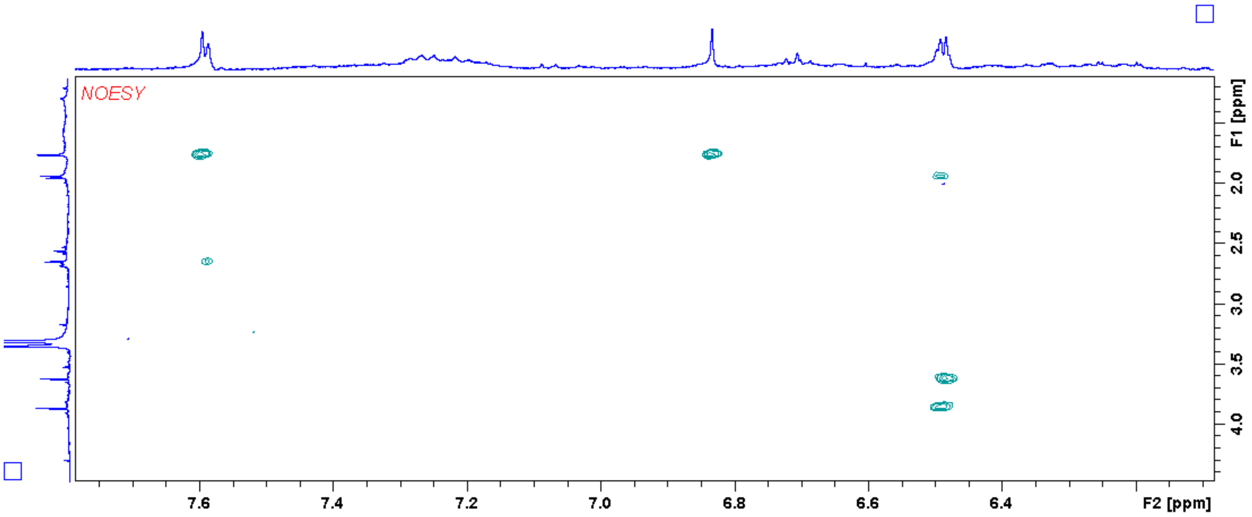

**COMPOUND 15 (fasamycin F)**

**Supplementary Figure 46 | Chemical structure of compound 13**

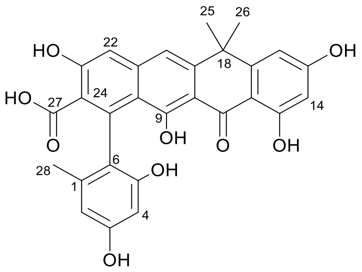


Molecular formula: $C_{28}H_{22}O_9$

Isolated yield: 3.4 mg

UV (PDA): $\lambda_{max}$ = 247, 287, 353, and 414 nm

HRMS (ESI) *m/z*: calculated $[M + H]^+$ = 503.1337; observed $[M + H]^+$ = 503.1326, Δ = -0.29 ppm.

**Supplementary Table 13 | NMR data for compound 13 in $CD_3OD$ at 400 MHz for $^1H$ and 100 MHz for $^{13}C$.**

| Position | $\delta_C$ ppm | $\delta_H$ ppm (no. of protons, multiplicity, J in Hz) | COSY | HMBC | NOESY |
|---|---|---|---|---|---|
| 1 | 139.56 | | | 28 | |
| 2 | 108.89 | 6.2 (1H, d, 2.30) | 4 | 4, 28 | 28 |
| 3 | 158.20 | | | | |
| 4 | 100.90 | 6.2 (1H, d, 2.30) | 2 | 2 | |
| 5 | 156.17 | | | 4 | |
| 6 | 121.28 | | | 2, 28 | |
| 7 | 138.31 | | | | |
| 8 | 117.68 | | | 20, 22 | |
| 9 | 167.74 | | | | |
| 10 | 108.24 | | | 20 | |
| 11 | 191.77 | | | | |
| 12 | 108.75 | | | 16 | |
| 13 | 166.89 | | | 14 | |
| 14 | 102.25 | 6.2 (1H, d, 2.22) | 16 | 16 | |
| 15 | 166.70 | | | 14, 16 | |

| | | | | | |
|---|---|---|---|---|---|
| 16 | 107.21 | 6.6 (1H, d, 2.22) | 14 | 14 | 25, 26 |
| 17 | 155.87 | | | 25, 26 | |
| 18 | 39.96 | | | 16, 20, 25, 26 | |
| 19 | 147.72 | | | 25, 26 | |
| 20 | 115.95 | 7.3 (1H, s) | | 22 | 25, 26 |
| 21 | 142.60 | | | | |
| 22 | 110.64 | 7.1 (1H, s) | | 20 | |
| 23 | 157.09 | | | 22 | |
| 24 | 128.39 | | | 22 | |
| 25 | 34.67 | 1.7 (3H, s) | | 26 | 16, 20 |
| 26 | 34.84 | 1.7 (3H, s) | | 25 | 16, 20 |
| 27 | 171.95 | | | | |
| 28 | 20.86 | 1.9 (3H, s) | | 2 | 2 |

**Supplementary Figure 47 | $^1H$ NMR spectrum ($CD_3OD$, 400 MHz) for compound 13.**

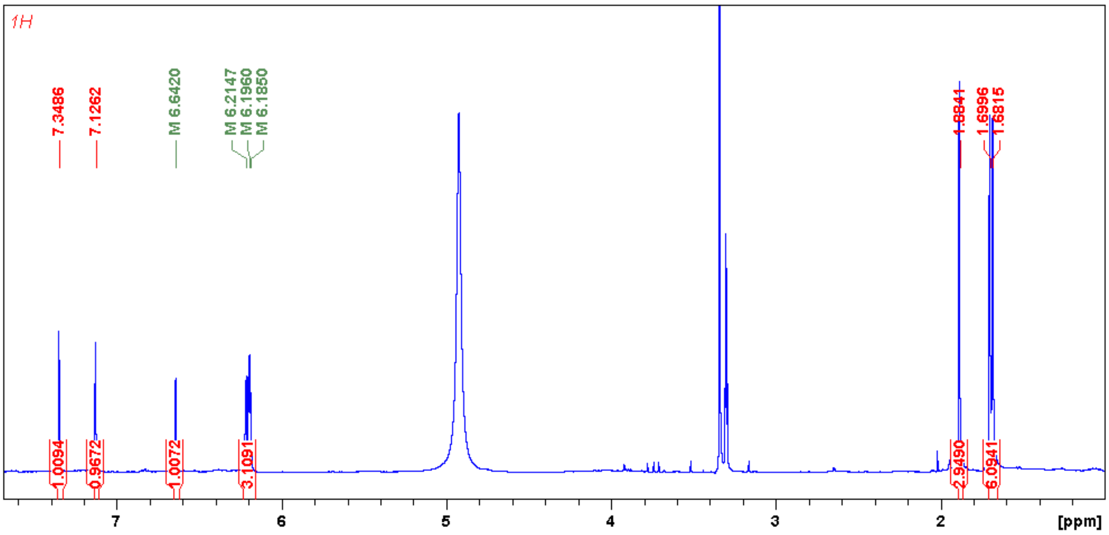

**Supplementary Figure 48 | $^{13}C$ NMR spectrum ($CD_3OD$, 100 MHz) for compound 13.**

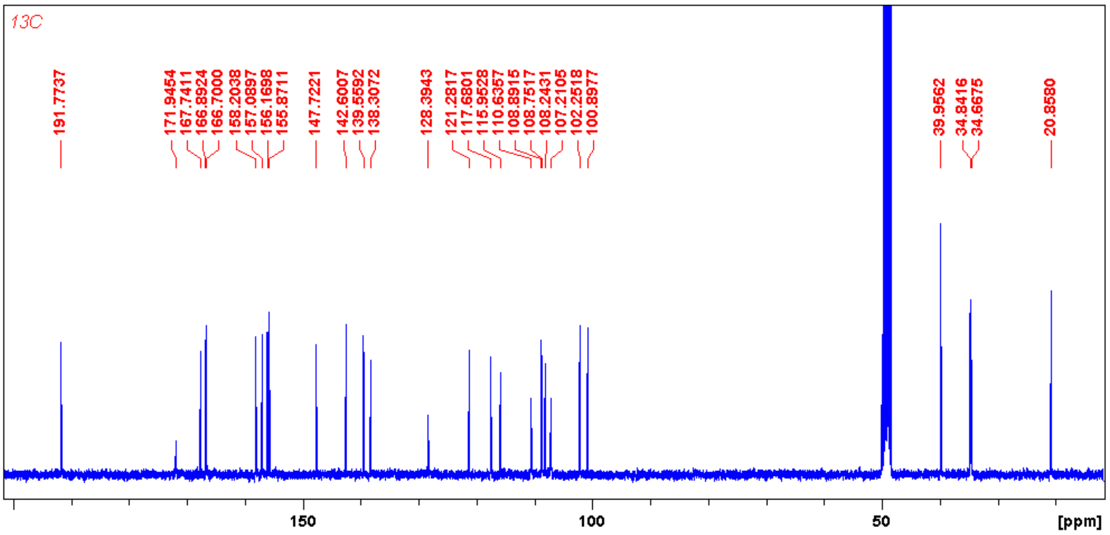


**Supplementary Figure 49 | HSQC spectrum ($CD_3OD$) for compound 13.**

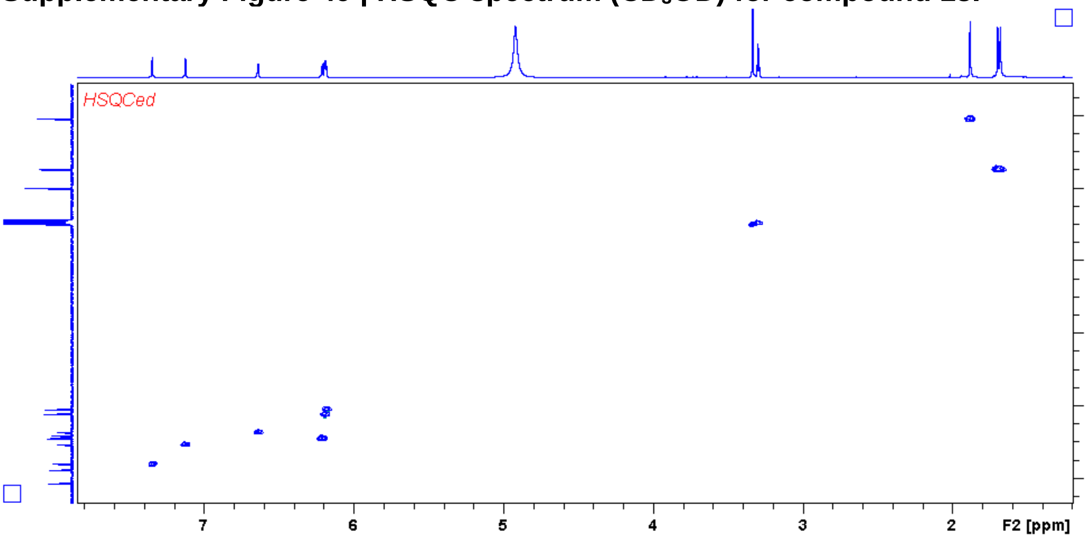


**Supplementary Figure 50 | HMBC spectrum ($CD_3OD$) for compound 13.**

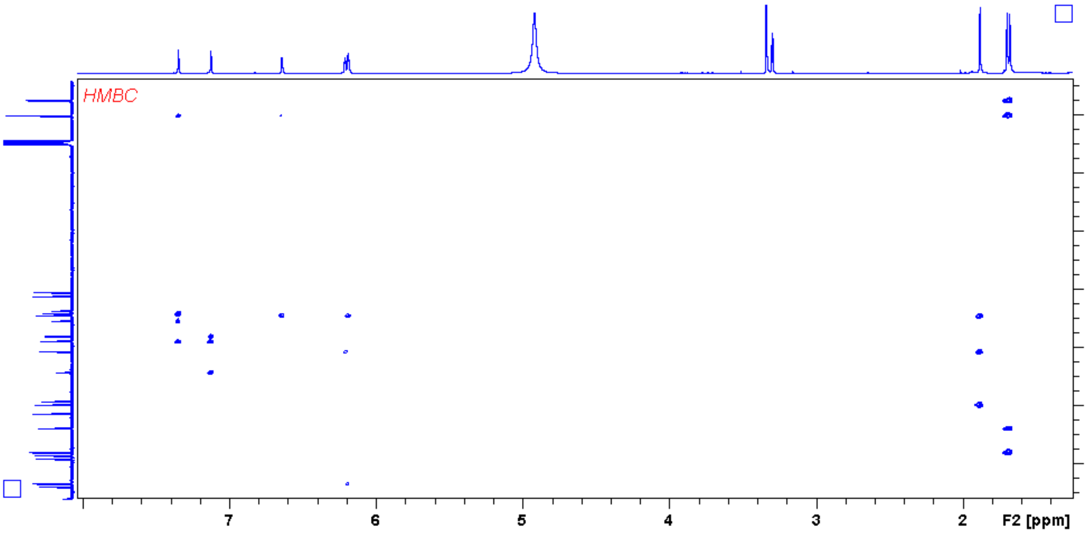

**Supplementary Figure 51 | NOESY spectrum ($CD_3OD$) for compound 13.**

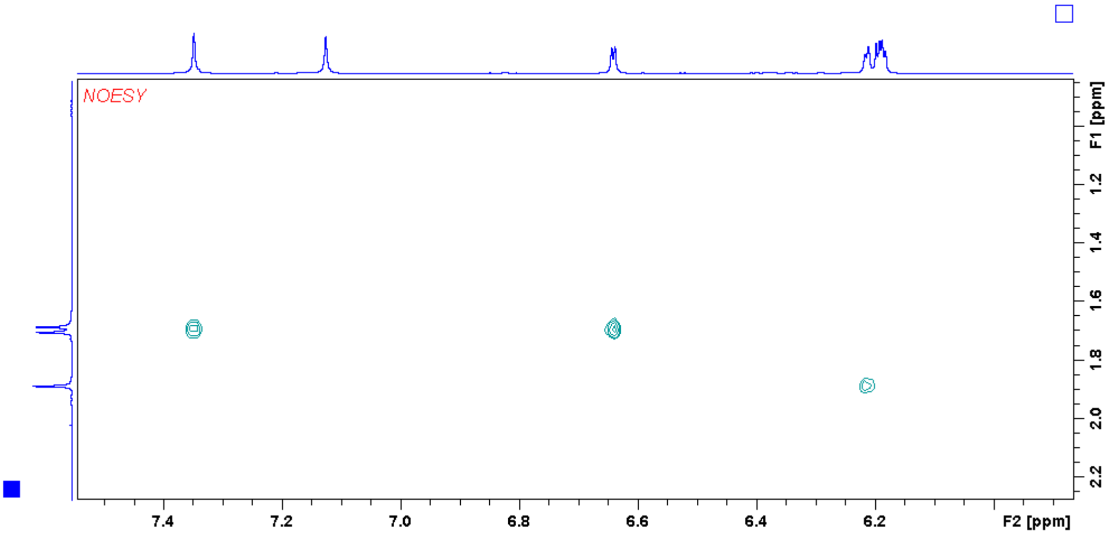
